# High throughput genotyping of structural variations in a complex plant genome using an original Affymetrix® Axiom® array

**DOI:** 10.1101/507756

**Authors:** Clément Mabire, Jorge Duarte, Aude Darracq, Ali Pirani, Hélène Rimbert, Delphine Madur, Valérie Combes, Clémentine Vitte, Sébastien Praud, Nathalie Rivière, Johann Joets, Jean-Philippe Pichon, Stéphane D. Nicolas

## Abstract

**Background:** Insertions/deletions (InDels) and more specifically presence/absence variations (PAVs) are pervasive in several species and have strong functional and phenotypic effect by removing or drastically modifying genes. Genotyping of such variants on large panels remains poorly addressed, while necessary for approaches such as association mapping or genomic selection.

**Results:** We have developed, as a proof of concept, a new high-throughput and affordable approach to genotype InDels. We first identified 141,000 InDels by aligning reads from the B73 line against the genome of three temperate maize inbred lines (F2, PH207, and C103) and reciprocally. Next, we designed an Affymetrix® Axiom® array to target these InDels, with a combination of probes selected at breakpoint sites (13%) or within the InDel sequence, either at polymorphic (25%) or non-polymorphic sites (63%) sites. The final array design is composed of 662,772 probes and targets 105,927 InDels, including PAVs ranging from 35bp to 129kbp. After Affymetrix® quality control, we successfully genotyped 86,648 polymorphic InDels (82% of all InDels interrogated by the array) on 445 maize DNA samples with 422,369 probes. Genotyping InDels using this approach produced a highly reliable dataset, with low genotyping error (~3%), high call rate (~98%), and high reproducibility (>95%). This reliability can be further increased by combining genotyping of several probes calling the same InDels (<0.1% error rate and >99.9% of call rate for 5 probes). This “proof of concept” tool was used to estimate the kinship matrix between 362 maize lines with 57,824 polymorphic InDels. This InDels kinship matrix was highly correlated with kinship estimated using SNPs from Illumina 50K SNP arrays.

**Conclusions:** We efficiently genotyped thousands of small to large InDels on a sizeable number of individuals using a new Affymetrix^®^ Axiom^®^ array. This powerful approach opens the way to studying the contribution of InDels to trait variation and heterosis in maize. The approach is easily extendable to other species and should contribute to decipher the biological impact of InDels at a larger scale.

## Background

In the past decade, there has been growing evidence that structural variations (SVs) are pervasive within plant genomes [1–9]. Insertion/deletions (InDels) are one class of SVs of particular interest, since they lead to the presence or absence of, sometimes large, genomic regions at a given locus, among individuals from the same species. The content of these InDels can be present elsewhere in the genome, but they can also be completely absent from the genome, in which case they are referred to as presence/absence variants (PAVs). Some InDels carry entire genes or affect gene regulatory elements and are thus likely to have a functional and phenotypic impact [10–12, 7, 13]. Hundreds to thousands of SVs, including PAVs and copy number variations (CNVs), have been discovered in several plant species, including wheat [14], rice [15], *Arabidopsis thaliana* [13], potato [16], pigeon peas [17], and sorghum [18]. These results support the idea that one single reference genome cannot properly represent the complete gene set of a given species. There has been an increasing interest for building new individual genomes in complement to the reference genome, in order to better describe the genetic diversity within a plant species [3, 19–25].

In maize, BAC sequence comparison first revealed that gene and transposable element content greatly vary between inbred lines [26, 27]. Whole genome sequencing of the B73 inbred line then provided the opportunity to explore the extent of SVs across the entire maize genome [28] by designing Comparative Genomic Hybridization (CGH) technology [29]. Several CGH studies found multiple CNVs between the B73 reference genome and other maize inbred lines or teosintes [2, 8, 9]. These studies demonstrated the large extent of SVs among maize inbred lines, including presence/absence variations of low copy sequences, such as genes. This was well illustrated by the discovery of a large 2 Mbp presence/absence region between Mo17 and B73 carrying several genes [2, 9, 20, 21]. However, CGH array technology shows several major drawbacks since (i) it does not allow the discovery of sequences that are not present in the reference genome used for designing probes of the arrays, (ii) it has a limited resolution which does not allow detection of InDels smaller than 1kb, and (iii) it is costly and labor-intensive, and therefore not adapted for genotyping several hundreds of individuals.

Methods based on SNP array experiments have been developed to detect CNVs and were shown to be more affordable and with higher throughput than CGH arrays [32–35]. Didion et al. (2012) identified atypical patterns of reduced hybridization intensities that were highly reproducible, so called “off-target variants” (OTVs) [36]. OTV patterns could originate either from the absence of the sequence due to a PAV polymorphism, or to a single nucleotide polymorphism within the probe sequence, thus preventing the correct hybridization of the DNA sample. For instance, 45,974 OTVs were discovered in a maize population using the 600K Affymetrix^®^ Axiom^®^ SNP array [37]. While these approaches proved to be useful, there is a strong risk of false positive detection of PAVs using OTV patterns, mainly because these arrays were not designed to target PAVs. In order to reduce this risk of false positive detection of PAVs and more largely CNVs, several methods based either on segmentation or Hidden Markov Chain have been developed to use variation of fluorescent intensity signal of contiguous probes along the genome [38–43] These kind of approaches have been used on 600K Affymetrix^®^ Axiom^®^ SNP array to detect several hundreds of CNVs and to explore the contribution of CNV to phenotypic variation [44].

With the emergence of massive parallel sequencing, new methods have been developed to detect structural variations based on the alignment of resequencing reads onto a high quality reference genome sequence. Among these, three have been mainly used [45] : (i) the “read-depth” (RD) method, which can only detect copy number variations; (ii) the “read-pair” (RP) method, which can detect deletions as well as small insertions (up to the size of the library insert); and (iii) the “split-read” (SR) method which can also detect deletions and small insertions (up to the size of a read). Chia et al. (2012) used the RD approach to identify CNVs among 104 maize lines and performed association studies for several traits [10]. However, the RD method does not allow the identification of novel sequences and is error prone, especially regarding the size of the discovered CNVs which greatly depends on the size of the sliding window used. The RP method has been implemented in many computational tools like BreakDancer [46] and has been widely used. Although it has proven to be highly efficient to detect deletions [47–49], this approach suffers from two limitations: it does not allow precise detection of breakpoints, and the size of the insertions which can be detected is directly limited by the library insert size. The SR method, which was first implemented in PInDel [50], has the advantage of defining breakpoints at a single-base resolution, but again the size of the detectable inserted sequence is limited.

The “assembly” (AS) method is able to detect all types of SVs of any size, but is also the most cost and computation-intensive. It is the only method able to detect large insertions with precise breakpoint definition. However, the assembly of large and complex genomes such as maize remains very expensive and computationally intensive, despite recent progress in this area [20, 21, 31]. There has been in the past some attempts to reduce this complexity by reducing the number of sequences to assemble. For instance, Lai et al., (2010) identified 104 deletions and 570 insertions among 6 maize inbred lines by assembling genomic regions from reads that did not map on the B73 reference genome [51]. The sequences assembled by this approach were enriched in erroneous reads or reads coming from external contamination, and they were too short to be anchored to the reference genome B73. Hirsch et al. (2014) identified several putatively expressed genes that were not present within B73 reference genome by assembling and comparing the transcriptome of hundreds of inbred lines [12]. This new approach was limited to the transcribed part of the genome and suffered from a high level of false positives. More recently, Lu et al., (2015) used genotyping by sequencing approaches on 14,129 inbred lines to identify 1.1 million short and unique sequences (GBS tags) that (i) did not align on the B73 reference genome, or were aligned but outside of a 10Mbp windows around their mapped position; or (ii) were mapped at the same location by joint linkage mapping in NAM populations using co-segregation with a SNP and logistic regression between the InDel and the SNP in an association panel [13]. The main drawback of this approach is the high percentage of missing data due to the low depth of sequencing, which requires imputation before being able to perform genetic analysis. Recent whole genome sequence assemblies of PH207 [31], and F2 [20] have allowed the identification of thousands of large InDel and PAV sequences. For instance, 2,500 genes were found either present or absent in PH207 and B73 genomes and 10,735 PAV sequences larger than 1kb were discovered between F2 and B73, including 417 novel genes in F2. These discovery approaches have been limited to a few individuals due to sequencing costs and computational challenges, so they have not been adapted for characterization of SVs on large maize panels. Darracq et al. (2018) developed an interesting approach for the genotyping of PAVs from mapping of low depth (5-20X) resequencing datasets [20]. This method is based on the comparison of reads aligning to the region found in F2 and in the line of interest. While this method is potentially adapted to genotype PAVs on any set of line with low resequencing data, it has been so far used for PAV genotyping on a low (<30) number of maize lines. Moreover, it is restricted to the analysis of PAVs, and is not adapted for genotyping other types of SVs. To avoid this ascertainment bias due to use of a single reference genome to genotype SV, other studies proposed to call SV by aligning reads on a pan-genome representing the combination of several genomes [14, 22, 52]. However, these approaches remained computationally challenging on a sizable set of individuals, time demanding, and costly for large and complex genomes, since it requires high-depth sequencing [52]. To our knowledge, no high-throughput genotyping approach has been developed for genotyping large numbers of InDels, including PAVs, on a large set of individuals. We have developed, as proof of concept, a new high-throughput and affordable array that is able to genotype simultaneously large insertions and deletions, with highly variable size and contents that are previously discovered by different sequencing methods. In this study, we present this approach which is both (i) comprehensive, as it includes the discovery and localization of deletions as well as insertions regarding the B73 reference genome at the base pair level and (ii) high-throughput, as it allows genotyping of thousands of InDels on hundreds of individuals. Our strategy takes advantage of next generation sequencing (NGS) technologies and recent advances in assembly of complex genomes. It also benefits from the high efficiency of SNP arrays like the high-throughput Affymetrix^®^ Axiom^®^ technology. In this paper, we detail how we discovered thousands of small to large InDels, including PAVs, from three maize inbred lines (F2, PH207 and C103) as compared to the B73 reference genome. We then describe how we designed and selected 600,000 probes to create a new Maize Affymetrix^®^ Axiom^®^ array to genotype these InDels. Finally, we describe how we successfully used this array to genotype an association panel of 362 maize inbred lines.

## Results

### InDel and PAV discovery

To design a comprehensive InDel genotyping array, we first discovered a set of InDels which would be representative of the maize temperate germplasm. We already had access to sequence data for the European flint line F2, and we benefited from a first set of 42,330 F2-specific sequences, larger than 150pb and totaling 16Mbp. This dataset was derived from the *de novo* assembly of an F2 paired-end that failed (at least for one read of the pair) to align onto the B73 AGPv2 sequence, and which were totally devoid of coverage by B73 reads (“Reference guided assembly” in Additional file 2: Figure S1, “no map” approach). We also took advantage of the work done by [20] to add another 10,044 F2-insertions (size >1 kb, total size of 88Mb), with less than 70% of their length covered by B73 reads discovered by a whole genome assembly approach (Additional file 2: Figure S1).

To complement these two datasets of F2/B73 deletions and insertions, we generated and assembled Illumina^®^ paired-end and mate-pair sequences from two other key founders of temperate maize breeding programs: PH207 and C103. We then used this F2, PH207, and C103 sequence data to detect all InDels, including PAVs, at base-pair resolution, between these three lines and B73. As opposed to the “reference guided assembly approach”, the “whole genome assembly” methodology allowed us to access both to their sequences and their breakpoints, permitting the genotyping of such InDels in several individuals (more details in Methods). We did not use the “no map” approach for InDel discovery on PH207 and C103, because this approach did not give access to breakpoint resolution, did not allow the discovery of InDels without knowledge of the specific sequence, and was almost redundant with the assembly approach.

We first aligned F2, PH207, and C103 sequences against the B73 reference genome sequence in order to detect deletions. Here, the term “deletion” does not reflect any underlying biological process of DNA excision but refers to a sequence of at least 100bp present in the B73 genome at one locus and absent in another line at the same locus. Deletions were detected for the three lines simultaneously using the “genotyping” option of PInDel [50], generating a set of 26,368 non-redundant deletions with precise identification of their breakpoints (Additional file 2: Figure S2A). The number of deletions found for each line was similar, respectively 12,165, 11,922, and 13,432 for F2, PH207, and C103. 67% of the deletions found were unique to one line, 24% were shared by two lines, and 9% by three lines (Additional file 2: Figure S2A). These results confirm the good complementarity of the lines chosen to discover InDels. The high proportion of unique deletions among 4 lines may also reflect that numerous InDels remain to be discovered in temperate maize germplasm.

Next, we generated a draft genome assembly for each of these lines, which was used as a template for alignment of B73 reads to detect insertions relative to the B73 reference genome (Additional file 1: Table S1). As for deletions, here the term “insertion” does not reflect any underlying biological process of DNA integration, but defines a sequence larger than 100bp that is present in one line at a given locus, and absent from B73 at the same locus. These three draft assemblies cover less than one third of the expected maize genome size but include a large portion of low copy sequences, including genes, as shown by BUSCO results (Table 1).

**Table 1:** F2, PH207, and C103 de novo assembly metrics.

| Maize line | Number of scaffolds | Min size | Max size | Average size | N50 | Total (Mb) | % of Ns | Complete BUSCOs (C) | Fragmented BUSCOs (F) | Missing BUSCOs (M) |
| --- | --- | --- | --- | --- | --- | --- | --- | --- | --- | --- |
| **F2** | 76,563 | 892 | 112,956 | 16,900 | 14,042 | 646.3 | 9.48% | 89.3% | 4.9% | 5.8% |
| **PH207** | 81,688 | 884 | 2,024,489 | 29,557 | 16,860 | 797.5 | 8.90% | 91.8% | 2.7% | 5.5% |
| **C103** | 84,990 | 886 | 120,582 | 19,305 | 16,146 | 793 | 8.21% | 90.6% | 4.2% | 5.2% |

Detection of insertions was processed separately for each inbred line and generated 28,221 insertions for F2, 27,904 insertions for C103, and 26,795 insertions for PH207, with their precise breakpoints (Additional file 2: Figure S2B). The number of insertions is similar between lines, but significantly greater than the observed deletions. Among these insertions, 26,691 cases could be uniquely anchored at base pair resolution onto the B73 reference genome sequence (Additional file 2: Figure S2B). Again, a majority of insertions were unique to one line (72%) confirming the complementarity of the material chosen (Figure S2B).

Finally, the results from the different approaches were merged into a non-redundant set of 141,325 InDel sequences (see Methods), comprising 52,175 deletions and 89,150 insertions. These regions were then used for the design of genotyping probes.

### Design of the genotyping array

#### Genotyping strategy

Large InDels can be efficiently genotyped with a SNP array using a combination of two types of probes: (i) “external” probes, which target breakpoints using the two flanking sequences of a given InDel (BP probes), and (ii) “internal” probes, which target presence/absence regions (PARs) within the internal sequence of InDels on polymorphic (OTV probes) or monomorphic sites (MONO probes). We define PARs as small portions of DNA sequence of at least 35bp that were observed present or absent at the genome level, when comparing two individuals. They are thus suitable for the design of presence/absence genotyping probes. Ideally, each InDel should be called by two BP probes on either side and by multiple internal probes, regularly distributed along the internal sequence of the InDel (Figure 1A). However, in practice, this combination of different probes is not always possible. For instance, precise breakpoints were not determined for all PAVs from our “no map” approach and [20], and PARs for internal probes were not always found in our InDels.

**Figure 1:**
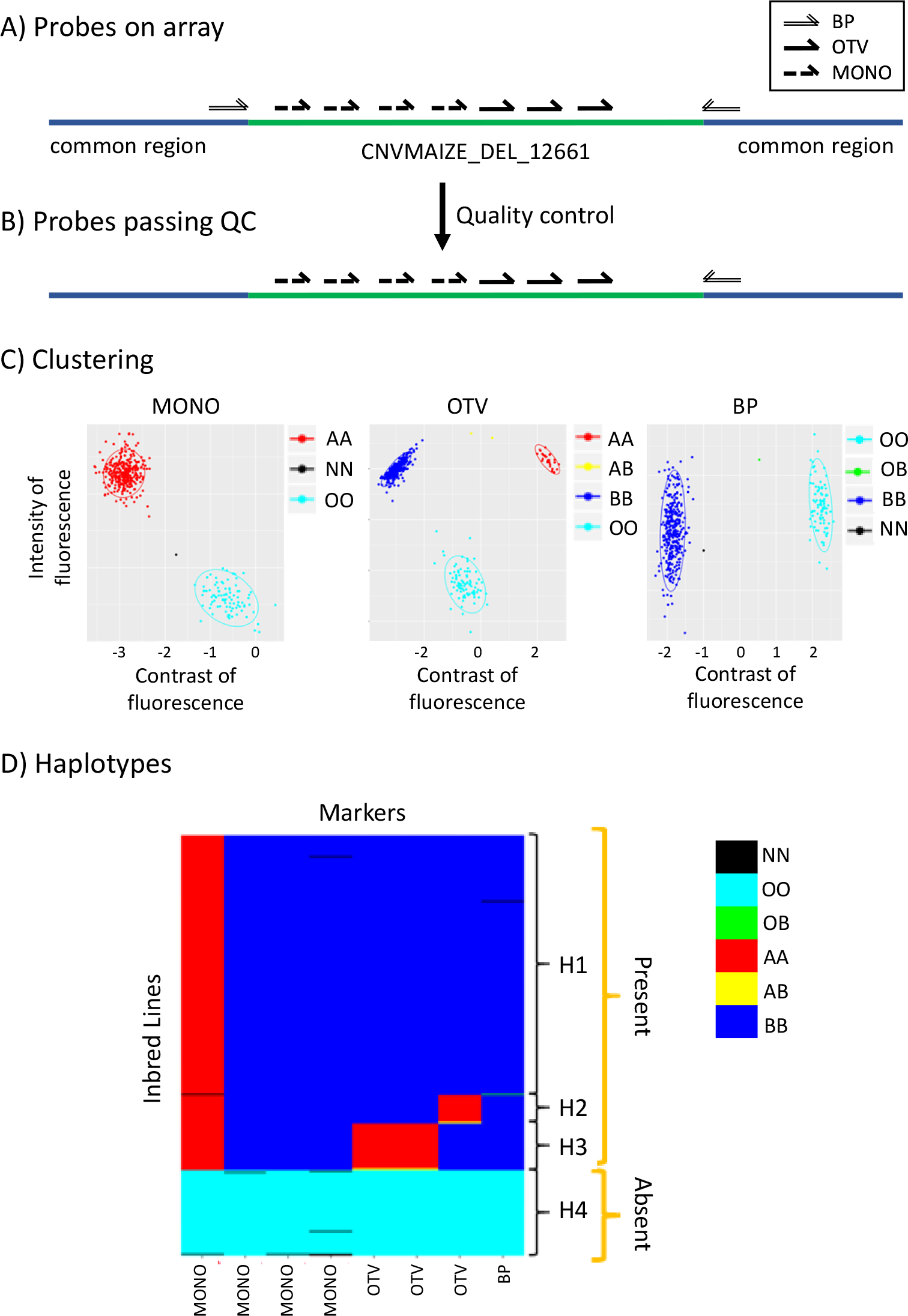
Genotyping of InDel CNVMAIZE_DEL_12661 using three probe types on 445 individuals. A) Schematic distribution of the 9 probes along the sequence of InDel CNVMAIZE_DEL_12661 (green line) and the bordering sequence common between all individuals (blue line) genotyped by the array. Double, dotted, and full arrows represented the probes designing on the forward and reverse flanking sequences of the breakpoint sites (BP), at not polymorphic (MONO) and polymorphic sites (OTV) within internal sequence of InDel. B) Schematic distribution of the 8 probes passing Affymetrix^®^ quality control and called by the Affymetrix^®^ pipeline C) Clustering produced by the Affymetrix^®^ algorithm for an OTV, MONO, and BP probe from InDel based on both fluorescence contrast (X axis) and intensity (Y axis) of the 445 inbred lines. Red, blue and yellow dots indicated the presence of the sequence (genotype “present”) either homozygous for allele A (AA) or allele B (BB) or heterozygous (AB), respectively. Cyan and green indicated that the sequence was absent in the individual (OO), or only in one copy of the sequence, e.g hemizygous for presence/absence (OB or OA). Black dots indicated individuals for which no genotype could be assigned (Missing data) D) Haplotypes displayed by the genotyping using 8 probes (column) on the 445 inbred lines (row). Colors corresponded to the genotype of individuals produced by clustering in C)

#### Probe design

BP probes should behave like classical SNP probes where one allele corresponds to the presence and the other to the absence of the InDel. They are useful to explore the conservation of the localization of large insertion/deletion events across multiple individuals, even when no internal probe can be designed due to the absence of PARs. Among the 141,325 selected variants, 86,406 InDels (22,420 deletions and 63,986 insertions as compared to the B73 reference genome sequence) had breakpoints defined at base-pair resolution and were suitable for BP probe design. Four different breakpoint types were identified according to the presence of micro-homology and/or shorter non homologous sequence [53] in place of a complete deleted sequence (Additional file 2: Figure S3): (type I) 3,397 cases with sharp breakpoints; (type II) 45,987 cases with a micro-homology sequence (8.6 bp on average and no more than 237 bp) which was present in one copy in the reference sequence and duplicated at both extremities of the novel inserted sequence; (type III) 36,893 cases harboring insertion of a short non-homologous fragment (42.2 bp on average and up to 892 bp) in place of a large deleted sequence; and (type IV) 156 cases with a combination of type II and type III breakpoints. Following Affymetrix^®^ recommendations, 19,010 InDels with type II breakpoints having a micro-homology sequence longer than 5bp were excluded from the design process. In the end, 67,396 InDels, representing 48% of all available InDel variants, were submitted to the Affymetrix^®^ design pipeline. Two probes, one on forward (FW) and one on reverse (REV) strand, were designed for each breakpoint. These probes were classified as *not possible* (18%), *not recommended* (33%), *neutral* (15%) and *recommended* (35%) by this automated pipeline (see Methods for details), leaving 33,430 InDels (51%) that could be targeted by at least one *recommended* probe.

Internal probes, which should behave like “off-target” variants [36] where the hybridization of the probe indicates presence of the InDel, and the absence of hybridization of the probe indicates absence of the InDel, are useful to explore the genetic diversity within InDel sequences (Figure 1 D). They will also be particularly interesting to target InDels for which no breakpoint could be identified (such as PAVs from the “no map” approach).

For the design of OTV probes, we benefited from the availability of SNPs which had been previously identified from the alignment of resequencing data from a core collection of 25 temperate maize inbred lines against the B73-F2 maize pan-genome from [20]. As a consequence, OTV probes have only been designed for deletions positioned on the B73 reference genome and F2 insertions coming from [20]. Among these, the context sequences of 436,162 SNPs, corresponding to 21,390 InDels, were extracted and submitted to the Affymetrix^®^ design pipeline. Two probes, one on forward (FW) and one on reverse (REV) strand, were designed for each SNP. A total of 872,324 OTV probes could be designed and scored as *not possible* (0.05%), *not recommended* (71%), *neutral* (14%) and *recommended* (16%), leaving 17,589 InDels (82%) which could be targeted by at least one *recommended* probe.

For the design of BP and OTV probes we could rely on Affymetrix^®^ design pipeline to identify probes localized in PARs and thus suitable for the Affymetrix^®^ Axiom^®^ technology. For the design of MONO probes, we first had to identify such PARs within 141,325 InDels cumulating 133Mbp of sequence. We used sequence masking methods to exclude repeats based on similarity to known maize repeats or on occurrence of 17-mers found within the sequencing datasets we had for B73, F2, PH207, and C103 (see Methods). By doing so, we identified 122,972 PARs, representing a cumulated size of 27Mbp, corresponding to 20.3% of the initial size and allowing the possibility to design MONO probes for 79,987 InDels (56.5%). These PAR sequences were successfully used for the design of 25,735,797 MONO probes, among which 59% were scored as *recommended* and allowed to target 62,875 InDels (79%).

With this combined approach, we designed a total of 26,715,361 probes targeting 117,756 InDels, which represent a cumulated length of 250 Mbp including 27 Mbp of PARs (Table 2).

**Table 2:** Number of probes and targeted InDels before and after selection for array design and passing the Affymetrix^®^ quality control according to different probes type. Percentages are indicated in brackets

|  | Before selection |  | On array |  | Called by Affymetrix® pipeline |  |
| --- | --- | --- | --- | --- | --- | --- |
|  | Probes | InDel<sup>+</sup> | Probes | InDel<sup>+</sup> | Probes | InDel<sup>+</sup> |
| **BP Type1** | 6,648<br>(0.02%) | 3,324<br>(2.82%) | 4,691<br>(0.71%) | 2,751<br>(2.6%) | 2,092<br>(0.44%) | 1,482<br>(1.66%) |
| **BP Type2** | 51,770<br>(0.2%) | 25,885<br>(21.98%) | 38,790<br>(5.85%) | 22,662<br>(21.39%) | 20,540<br>(4.29%) | 14,407<br>(16.12%) |
| **BP Type3** | 71,820<br>(0.27%) | 35,910<br>(30.5%) | 41,272<br>(6.23%) | 27,897<br>(26.34%) | 23,631<br>(4.93%) | 18,485<br>(20.68%) |
| **BP Type4** | 312<br>(0.001%) | 156<br>(0.13%) | 241<br>(0.04%) | 146<br>(0.14%) | 119<br>(0.02%) | 93<br>(0.1%) |
| **OTV** | 872,324<br>(3.26%) | 21,390<br>(18.16%) | 163,278<br>(24.64%) | 18,558<br>(17.52%) | 96,867<br>(20.22%) | 15,064<br>(16.85%) |
| **MONO** | 25,735,797<br>(96.25%) | 68,573<br>(58.23%) | 414,500<br>(62.54%) | 65,796<br>(62.11%) | 335,778<br>(70.1%) | 63,597<br>(71.14%) |
| **ALL** | 26,738,671 | 117,756 | 662,772 | 105,927 | 479,027 | 89,393 |

<sup>+</sup>Note that a same InDel could be genotyped by several probe types which resulted in the percentage values great than 1.

Among these InDels, 97,748 (83%) can only be targeted with either internal or external probes, but not both (Figure 3A). These results support our overall strategy which includes the discovery of InDels, with precise breakpoints in a preliminary step, and the use of complementary internal/external probes for the genotyping of large InDels.

**Figure 2:**
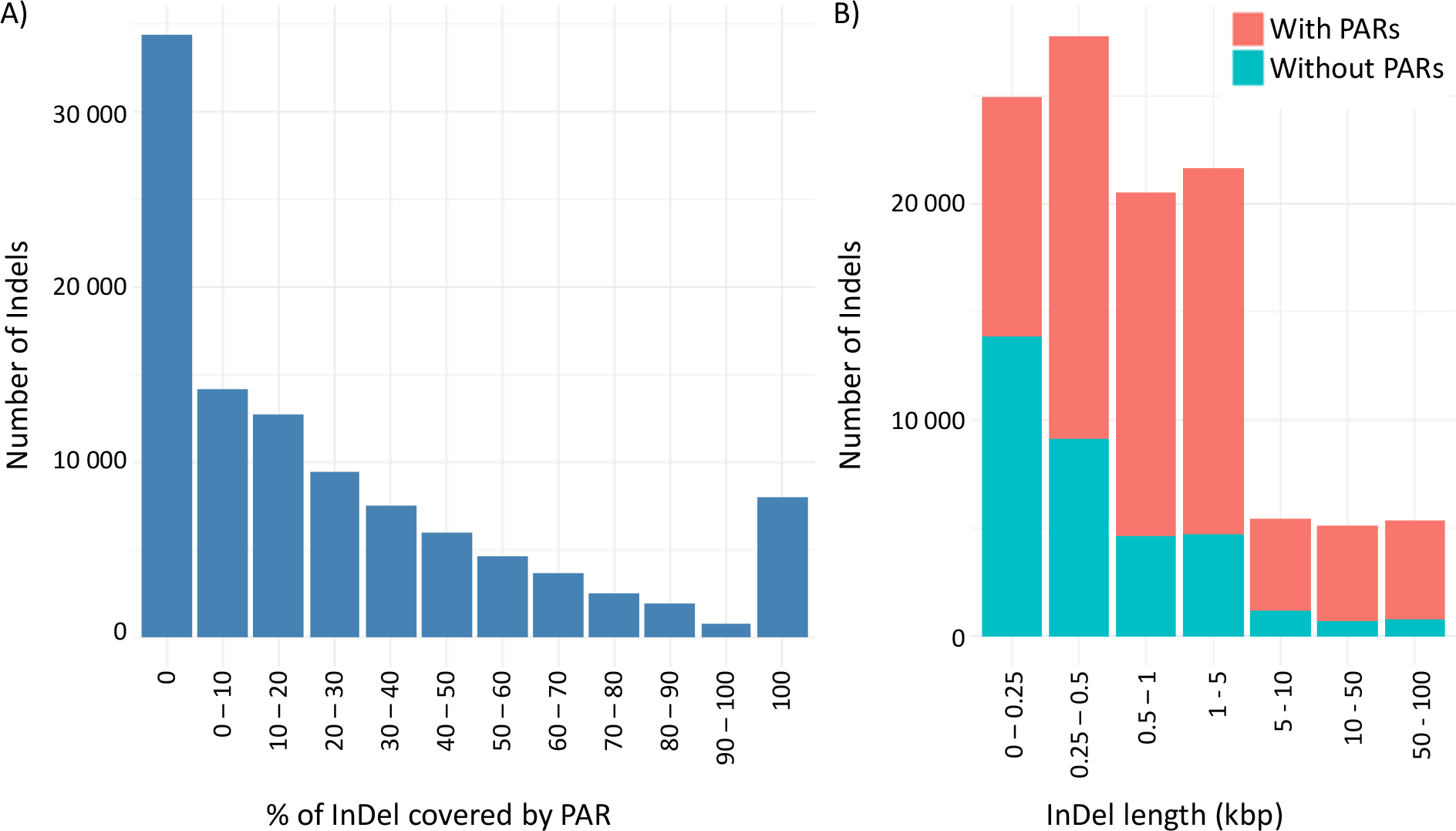
Distribution of 105,927 InDels genotyped by the array according to their size and the cumulated length of Presence/Absence regions (PARs) in their internal sequence. A) Distribution of the number of InDels according to the proportion of presence/absence regions (sequence not present elsewhere in the genome) within their internal sequence. B) Distribution of the number of InDels according to their size (kbp) and the percentage of internal sequence of InDel covered by PAR(s). Red Color indicates the proportion of InDels with (red) or without (blue) PARs for the 7 InDel size classes.

**Figure 3:**
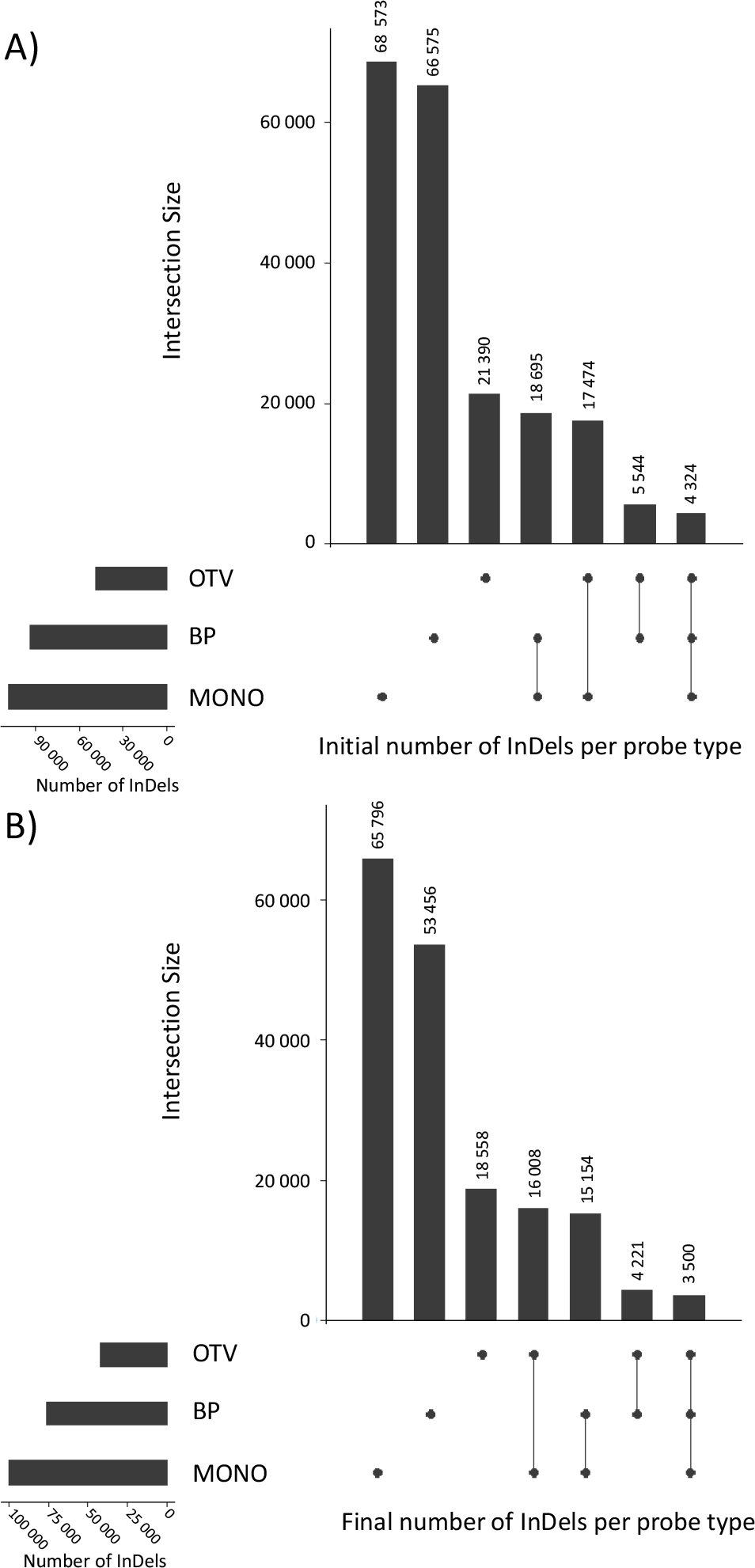
Number of InDels interrogated by each probe types or their combination, for which: a probe could be designed (A) and a probe was finally selected to be included in the final array (B). Vertical bars indicate number of InDels interrogated by each probe types or their combination. Black points and connected traits below the vertical bars indicate the corresponding probes types or their combination that are used for interrogating this subset of InDels. Horizontal bars indicate number of InDels interrogated by each probe types (OTV, BP, MONO). Number of InDels by probe type, for which: a probe could be designed (A) and a probe was finally selected to be included in the final array (B). Number of InDels that could be targeted by each type of probes designed (A) and selected to be included in the final array (B).

#### Array design

We used the Affymetrix^®^ recommendations to select the 700,000 probes to be included in the final array, plus some other criteria depending on the probe type. Nevertheless, because of their added value, we decided to keep all BP probes as long as they had less than 3 hits on the B73 reference genome sequence. This first selection consumed 84,994 probes targeting 53,456 InDels, among which 70% could only be targeted by BP probes. Concerning OTV and MONO probes, we first selected *neutral* and *recommended* probes having no hit at all (for insertions), and only one hit (for deletions), against the B73 reference genome sequence. We then considered their density with the objective to maximize the number of InDels that could be surveyed, as well as to have an even distribution of probes along targeted InDel sequences (see Methods). We then performed a second selection among *not recommended* OTV and MONO probes for 4,541 InDels that were still not targeted. After filtering some duplicated probes, we built a final array design containing 662,772 probes targeting 105,927 InDels that represent a cumulated length of 232 Mbp, including 25.9 Mbp of PARs.

#### Description of the array content

The final array design allows genotyping InDels with various sizes, ranging from 37 bp to 129.7 kbp, with a median of 501 bp (Figure 2). They are covered by 1 to 482 probes, with a median of 3 probes per InDel (Additional file 2: Figure S4). The number of probes does not always reflect the length of the InDels, as the proportion of PARs within InDels is highly variable (Figure 2A). 8,040 InDels (ranging from 37 bp to 2,409 bp, with a median of 163 bp) were completely covered by PARs and could thus be considered as a proper PAVs, 34,372 InDels (ranging from 101 to 129,700 bp with a median of 320 bp) were not covered by any PAR at all (Figure 2A). The biggest InDels contains more frequently PARs than the little ones (Figure 2B). In fact, the number of internal probes were more strongly correlated to the size of the PARs (r2 = 0.79) rather than to the size of the InDels (r2 = 0.16) (Additional file 2: Figure S5).

As expected, the probe selection process did not impact the overall distribution of probe types among targeted InDels, as 35% of them can exclusively be genotyped by BP probes, and 50% can only be genotyped by internal probes, among which 73% are only targeted by the use of the original MONO probes (Figure 3B). Indeed, a large number of InDels did not contain PARs and cannot be genotyped with 35bp internal probes but only with BP probes. Whereas, others InDels contains PARs but have no BP probes due to the InDel discovery approach (“no map”).

Among the 43,117 InDels that could be anchored onto the B73 reference genome sequence and which were included in the array design, 13,737 were located inside a gene, 57 close to a gene (less than 1 kb away), 1,311 inside a pseudo-gene and 2,212 inside a transposable element. InDels and probe density varied across each chromosome (Additional file 2: Figure S6). We observed a higher density in chromosome arms than in peri-centromeric regions (Additional file 2: Figure S6). We also identified clusters of InDels with a large specific sequence at the beginning of chromosome 6 (10-20Mbp) or at the end of chromosome 5 (~190Mbp).

### Assessing array quality by genotyping 105,927 InDels on 480 maize DNA samples

#### InDel calling using dedicated Affymetrix® pipelines

We genotyped 480 maize DNA samples including 440 inbred lines, 24 highly recombinant inbred lines and 16 F1 hybrids. Dedicated Affymetrix^®^ pipelines were implemented for each of the probe types to call genotype of the InDels based on fluorescent intensity and contrast variation of the probes. It included two algorithms already developed by Affymetrix^®^ [36] for BP and OTV probes (Additional file 2: Figure S7A et B) and a third one, which was newly developed for the calling of presence/absence genotypes using MONO probes (Additional file 2: Figure S7C). 35 DNA samples including all F1 hybrids, did not pass Affymetrix^®^ quality control due to their low call rate (<0.9) and were eliminated. Call rate of the 445 remaining samples, which are all inbred lines, varied from 96% to 99% with a median of 98%. The call rate varied according to probe type (median of 90% and 99% for BP and internal probes, respectively). Out of 662,772 probes, 479,027 probes representing 89,393 InDels passed Affymetrix^®^ quality control and were called on 445 DNA samples. Respectively 55%, 59%, and 81% of BP, OTV, and MONO probes were converted into recommended markers after clustering by Affymetrix^®^ pipelines (Additional file 1: Table S2, S3, and S4). 94% of these recommended BP and OTV markers were classified as “PolyHighResolution” (PHR) indicating a high quality of clustering and that these markers were polymorphic (Additional file 2: Figure S8). Note that the criteria defining high quality of clustering for MONO probes called by new Hom2OTV algorithm was not yet implemented in Affymetrix pipeline (Additional file 1: Table S4 and Additional file 2: Figure S7C). As a consequence, classification of MONO probes could not be comparable to BP and OTV probes. Thanks to the 3 probe types and redundancy, 84% of all InDels could be called with an average of 5.4 probes per InDel.

To evaluate the genotyping ability of the 479,027 probes, we first compared the clustering of inbred lines expected for three probe types (BP, OTV, and MONO) with the observed clustering of inbred lines based on fluorescence intensity and contrast of 445 inbred lines genotyped with the array. For BP probes, we expected at least two clusters corresponding to the individuals homozygous either for presence (“AA” or “BB”) or absence (“OO”). A third cluster could be observed when individuals were heterozygous individuals for presence/absence (“OA” or “OB” hemizygous) (Figure 1C). For OTV probes, we expected at least 3 different clusters: two cluster corresponding to the individuals homozygous for allele A or B of SNP (“AA”, “BB”), and a third “off-target” cluster for the individuals homozygous for absence (“OO”). A fourth cluster could be observed when some individuals were heterozygous at the within-InDel SNPs (AB). For MONO probes, we expected only two clusters corresponding to the individuals for which the sequence was present (“AA” or “BB”) or absent (“OO“, “AA” or “BB”) (Figure 1C). The observed clustering by the three dedicated pipelines was consistent with the expected clustering for 43% of BP, 83% of OTV and 63% of MONO probes (Table 3).

**Table 3:**
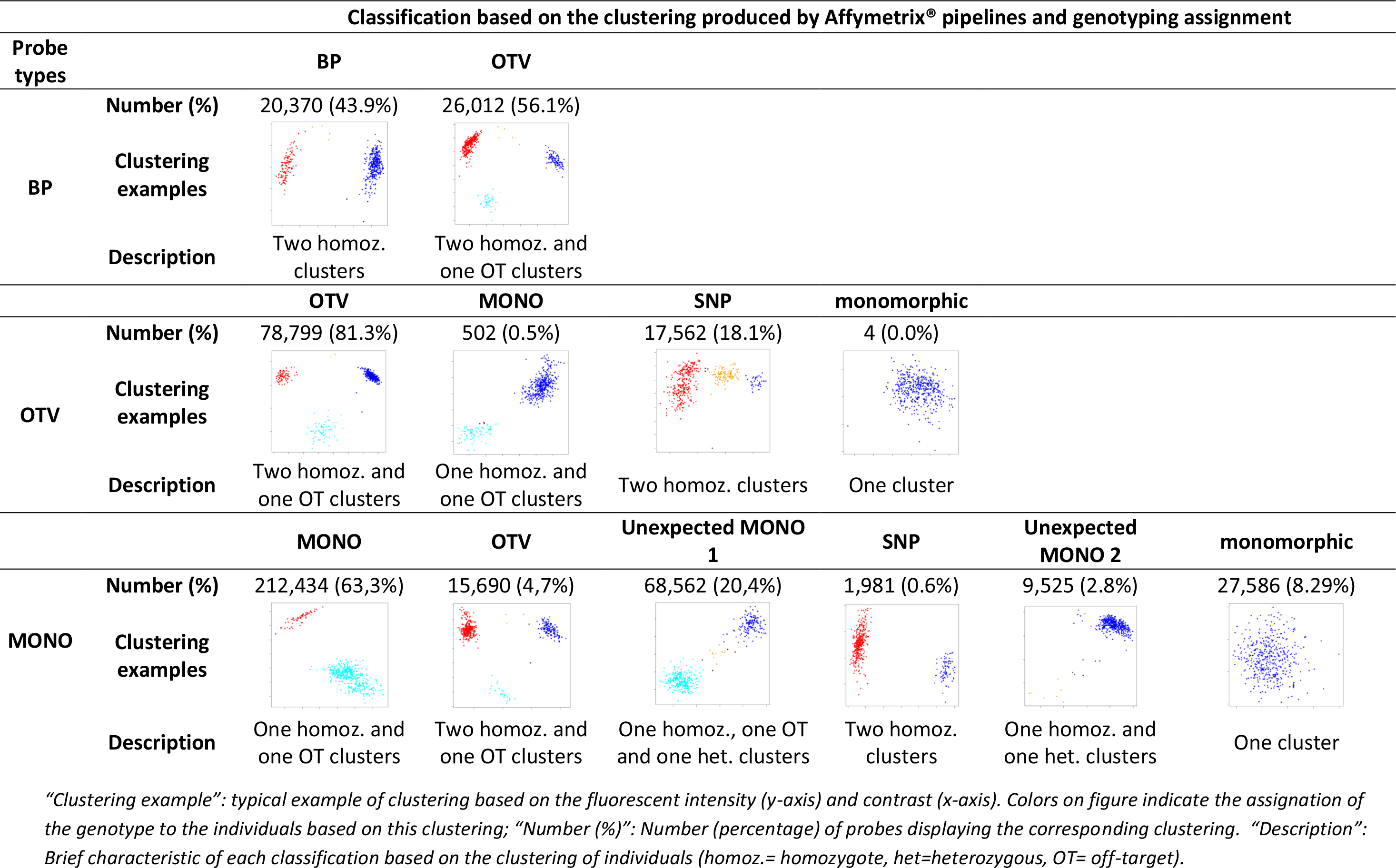
Comparison between the clustering expected for BP, MONO, and OTV probe types and the clustering produced by Affymetrix^®^ pipelines based on the fluorescent intensity and contrast of 445 inbred lines for 479,027 probes.

We observed also some unexpected clustering. For 57% of BP probes, we observed an additional off-target cluster (OTV in Table 3). This indicates that some BP probes did not hybridize properly in some inbred lines, which can either be due to the presence of polymorphism within flanking sequences of the targeted InDels or to the existence of more complex rearrangements removing the breakpoints.

Regarding MONO probes, 25% displayed additional cluster(s) when the sequence was present suggesting the presence of single nucleotide polymorphisms at this position. Among these, we were able to distinguish two types of clustering (Table 3). 4.7% of MONO probes exhibited a clustering similar to those observed for OTV probes suggesting that these MONO probes revealed, by chance, a single nucleotide polymorphism. In contrast, 20.4% of MONO probes displayed an unexpected clustering pattern for inbred lines with the presence of a heterozygous cluster but absence of a second homozygous cluster for SNP (Additional file 2: Figure S9B). In the end, 2.8% of MONO probes displayed an additional heterozygous cluster for SNP when the sequence is present but no “off target” cluster corresponding to individuals for which the sequence is absent (Additional file 2: Figure S9D).

For 18% of OTV (Additional file 2: Figure S9A) and 8.3% of MONO probes, clustering displayed no “off target” cluster for absence, suggesting no presence/absence polymorphism at this position (Table 3). Note that some BP were also classified as monomorphic for presence/absence but were filtered out by the BP pipeline (“MonoHighResolution” in Additional file 1: Table S2 and Additional file 2: Figure S8). These monomorphic probes originated from false positive discovery of InDels or PARs within InDels that are not present/absent elsewhere in the genome of four lines (see Discussion). After removing these monomorphic probes for presence/absence, 422,369 probes allowed us to successfully genotype a total of 86,648 InDels (82% of 105,927 InDels targeted by the array) on 445 inbred lines.

### Evaluation of genotyping reproducibility and quality

#### Consistency of genotyping among the four inbred lines used for InDel discovery

We used the 479,027 probes passing Affymetrix^®^ quality controls to evaluate the quality of Presence/Absence genotyping by comparing the genotyping results obtained from our array (GBA: Genotyping By Array) with those from sequencing (GBS: Genotyping by Sequencing) for the 4 lines used for the discovery of InDels (B73, F2, PH207, and C103). Respectively, 97%, 912%, and 88% of the BP, OTV, and MONO probes had a genotyping result consistent with results obtained from BLAST alignments against our three draft genome assemblies and the B73 reference genome. We observed a strong asymmetry for concordance rates for internal probes (OTV and MONO) depending on whether the genotype has been called by sequencing as present or absent (95% vs 80% present and absent, respectively, Table 4). Interestingly, we observed no asymmetry for BP probes that are designed exclusively on B73 genome compared to OTV and MONO probes that are designed from the 4 genome assemblies (Table 4). These low consistencies for internal probes when genotype by sequencing indicated absence could be explained by the use of incompletely assembled genomes of the three lines (PH207, C103, F2) to call the presence/absence genotype from sequencing.

**Table 4:** Consistency rate between genotyping by sequencing and by array for the 4 individuals used to discover the InDels, for the three probe types and for the two different genotypes observed from sequencing: presence (P) or absence (A).

| Probe Types | Genotype by sequencing | B73 | F2 | C103 | PH207 | All Lines |
| --- | --- | --- | --- | --- | --- | --- |
| BP | A | 0,98 | 0,98 | 0,98 | 0,97 | 0,98 |
|  | P | 0,97 | 0,97 | 0,97 | 0,96 | 0,97 |
|  | ALL | 0,97 | 0,97 | 0,97 | 0,97 | 0,97 |
| OTV | A | 0,85 | 0,89 | 0,80 | 0,78 | 0,83 |
|  | P | 0,93 | 0,97 | 0,96 | 0,96 | 0,96 |
|  | ALL | 0,90 | 0,95 | 0,91 | 0,90 | 0,92 |
| MONO | A | 0,77 | 0,81 | 0,82 | 0,81 | 0,80 |
|  | P | 0,90 | 0,98 | 0,94 | 0,94 | 0,95 |
|  | ALL | 0,82 | 0,94 | 0,89 | 0,88 | 0,88 |
| ALL | A | 0,80 | 0,86 | 0,84 | 0,82 | 0,82 |
|  | P | 0,92 | 0,97 | 0,94 | 0,95 | 0,95 |
|  | ALL | 0,85 | 0,95 | 0,90 | 0,89 | 0,90 |

*\*Note that consistency rate of hemizygous genotypes (heterozygous for presence / absence) were not displayed in the table for BP probes but considered to estimate global consistency rate (ALL). Note that the absence of probe sequence due to absence of hybridization or no alignment on draft sequence of BP probes were considered as missing data. Missing data were not included in the comparison for all probes.*

If the genomic region containing the InDels were absent or badly assembled in at least one line, some probes would not align properly, resulting in false absence calls, instead of presence in GBS. The four inbred lines showed very similar concordance rates, F2 being the most concordant (95%). This could be partially explained by the higher proportion of GBS present calls in F2 as compared to the three other lines since GBS present calls are more consistent with GBA than GBS absent calls. The median consistency rate of probes within InDels remained relatively high and stable, around 90%, independently of the number of probes per InDel (Figure S10), suggesting no relationship between the consistency rate of individual probes and length of PARs within InDels.

#### Consistency among probes from the same InDel

To estimate the consistency of different probes for typing a given InDel, we analyzed genotyping results for 50,648 InDels genotyped with at least two probes in a collection of 362 temperate inbred lines. For each InDel and each inbred line, we calculated the average allelic frequency of presence over all probes. Frequencies of 1 (presence) and 0 (absence) indicated that all probes displayed consistent genotyping for the corresponding inbred line (Figure 1D and Figure S11A). Alternatively, frequencies different from 0 or 1 (FreqDiff01) indicated that at least one probe displayed inconsistent genotyping with other probes for corresponding inbred lines (Figure S11B). Overall, 75% of the InDel genotyping resulted in an average allelic frequency for the presence of 1 or 0, meaning that all probes had a consistent genotyping results for calling the allele at both present or absent states, respectively (Figure 4A).

**Figure 4:**
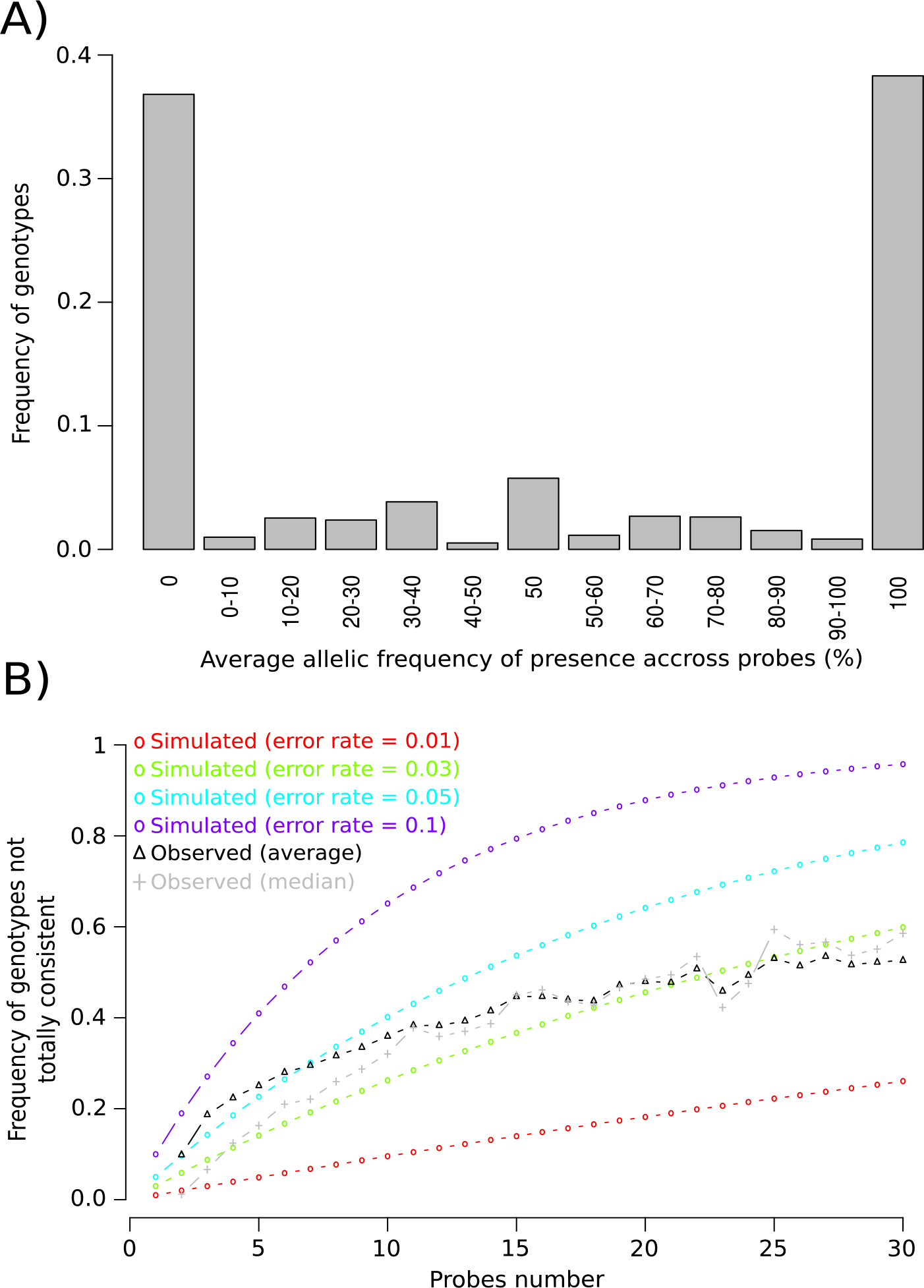
Consistencies among probes within 50,648 InDels with at least two probes genotyped in 362 inbred lines. (A) Distribution of the average allelic frequencies of present calls over all probes. (B) Variation of proportion of genotypes not fully consistent across all probes. The black and gray curves with triangle points represent the variation of the median and average FreqDiff01 across InDels, respectively. Colored curves with circle points represent the expected variation of the proportion for different error rates (1%: red, 3%: green, 5%: light blue, 10%: dark blue). Frequencies of 1 (presence) and 0 (absence) indicate that all probes had consistent genotypes for the corresponding inbred line. Intermediate frequencies (FreqDiff01) indicate that at least one probe was not consistent with the other probes for the same InDel in one inbred line.

However, we observed a strong variation of median (average) allelic frequency difference from 0 or 1 (FreqDiff01), according to the number of probe interrogating that InDel (Figure 4B, Additional file 1: Table S5). Median (average) FreqDiff01 across InDels varied from of 1.2% (9.8%) to 58% (52%) when the number of probes varied from 0 to 30 (Figure 4B, Additional file 1: Table S5). We compared this variation to what could be expected for different probe genotyping error rates (1%, 3%, 5%, and 10%). Based on this comparison, we estimated the probe genotyping error rate is approximately 3% (Figure 4). For InDels with fewer probes (<10), the average and median FreqDiff01 differed slightly, suggesting that some InDels with low probe numbers displayed high genotyping inconsistencies among their probes (Figure 4, Additional file 1: Table S5). In order to evaluate whether probe genotyping error is similar for present or absent calls, we analyzed the variation of FreqDiff01 with regard to the average frequency of absence of InDel sequences in 362 lines (Additional file 2: Figure S12A). The median FreqDiff01 was higher for InDels which have their sequence more frequently absent than present across 362 lines, regardless of the number of probes (Additional file 2: Figure S12B). It suggested that genotyping was more accurate for absence than presence. This was logical, considering that polymorphisms within probes would preclude hybridization of the probes for some lines, and it would result in absent calls with MONO and OTV probes, while polymorphisms within probes have no impact when the sequences are absent.

Combining genotyping from multiple probes within InDels greatly improved reliability of InDel genotyping, since it allowed (i) to correct the individual genotyping errors due to polymorphisms within probe sequences, (ii) to reduce the missing data rate due to bad clustering or probes polymorphisms, and (iii) to remove probes displaying highly-divergent genotypes compared to other probes for the same InDel, due, for example, to a bad design of the probes. In order to evaluate the combining of genotypes of several probes on the accuracy of InDel genotyping, we simulated global genotyping error rates for InDels by assigning to each inbred line the most frequent allele, based on the average frequency over all probes from an InDel, with various genotyping error rates (Additional file 1: Table S6). By this approach, the genotyping error for InDels was greatly reduced. Considering a probe genotyping error of 5%, the genotyping error of InDels for inbred lines were reduced to 0.2% and 0.1%, when the number of probes within the InDels were 2 and 5, respectively (Additional file 1: Table S6). Combining genotypes from several probes also strongly reduced the average missing data rate for InDels; it decreased from 2.3% to 0.2%, when the number of probes increased from 2 to 5 (Additional file 1: Table S5). However, some contradictory probe genotypes were repeatedly found across the 362 samples (Additional file 2: Figure S11B), suggesting that some probe inconsistencies could have biological origins (*i.e* more complex rearrangement), rather than being genotyping errors. Additionally, 35% of InDels called by BP had their FW and REV probes classified differently (e.g. one as BP and the other as OTV). Altogether, these results suggest that some calling inconsistencies between probes within InDels could come from polymorphisms in the flanking sequence while some other could be due to local rearrangements in the genotyped lines as compared to the lines used for InDels discovery.

#### Reproducibility and Mendelian inheritance

Genotyping reproducibility was evaluated by comparing genotypes between five DNA replicates corresponding to unique F1 hybrids derived from a cross between B73 and F72 for all probes type. Median reproducibility was 95%, 96%, and 97% for BP, OTV and MONO probes respectively. Interestingly, there is some variation of reproducibility relative to probe clustering (Additional file 1: Table S7). Note that Affymetrix^©^ algorithms were not specified to genotype hemizygote using OTV and MONO probes in this dataset. We also performed a parent-offspring analysis on 12 F1 hybrids derived from 9 parental lines by comparing genotypes observed of these F1 hybrids with those predicted from genotypes of their two parental lines for 46,382 BP probes (Additional file 1: Table S8). On average, 95% and 77% of observed genotypes were consistent with those predicted from parental lines for homozygous and hemizygous genotypes, respectively (Additional file 1: Table S8). The consistency rate was slightly higher when genotypes were absent (98%) than present (94.5%). Note that the seed-lot of parental lines used for producing F1 hybrids were different from those genotyped, which could explain lower consistencies rate than for DNA replicate of F1 hybrids. Note also that the genotypes of all F1 hybrids have been initially eliminated by Affymetrix^®^ quality control due to their low call rate and were therefore forced for reproducibility analysis. This low call rates can be attributed to the fact that these samples had different genotype cluster properties (probe intensity profiles) compared to the samples that passed QC. As a consequence, this strongly increased the missing data rate for the F1 hybrids for OTV and MONO probes.

In the end, we evaluated genotyping reproducibility for inbred lines, by comparing the genotyping results of 13 different inbred lines that were replicated in the experiment (Additional file 1: Table S9). Note that these are not perfect biological replicates, as they represent the same variety but come either from different seed lots or from different accessions. These replicates exhibited a genotyping difference varying from 0.6% to 5.2% (Median = 1.7%, Additional file 1: Table S9). This is similar to the amount of inconsistencies obtained on the same material using a 50K SNP array [54], suggesting that InDel genotyping inconsistencies for replicates can be attributed mostly to seed-lot divergences, rather than genotyping errors (Additional file 1: Table S9). However, genotyping reproducibility was higher for these inbred lines than for the DNA replicates of the F1 hybrid, suggesting that errors in F1 hybrids can mostly be attributed to the inability to genotype hemizygous with OTV and MONO probe for this small dataset.

### Application: Diversity analysis of 362 maize inbred lines panel

In order to evaluate this new array for genetic analysis, we analyzed genetic diversity using 57,824 polymorphic InDels on a subset of 362 inbred lines, representing genetic variation that has been successfully used to decipher maize genetic structuration and perform genome-wide association studies [55–57]. To represent each InDel in the diversity analysis, we selected one single probe per InDel, based on the probe genotyping quality (see Methods).

We first compared kinship values between 362 inbred lines estimated with 57,824 InDels and with 28,143 prefixed Panzea SNPs from the 50K SNP array. Kinship values between lines obtained with SNPs and InDels were strongly similar and highly correlated (r=0.9), except those for a couple of lines closely related to B73 and F2 (Additional file 2: Figure S13). Then, we performed Principal Coordinate Analysis (PCoA) based on the genetic distance between 362 lines estimated by InDels and SNPs (Figure 5). We included on this PCoA the genetic structuration of these 362 inbred lines, as obtained from the prefixed Panzea SNPs from the 50K SNP array [55]. The global genetic structure developed using two types of polymorphisms are highly similar. The first axis showed good discrimination of European Flint from Corn Belt Dent and Stiff Stalk lines, while the second axis discriminated European Flint and Northern Flint lines. Overall, the clustering of individuals based on genetic distance estimated with InDels (Figure 5A) was consistent with those estimated with SNPs (Figure 5B). We observed that B73 and F2, which were used to discover the majority of InDels, were more contrasted on PCoA when genetic distance was estimated with InDels, as compared with SNPs from the 50K array, indicating some ascertainment bias. We thus performed two PCoAs, with InDels and SNPs, excluding B73 and F2 (Additional file 2: Figure S14). The two PCoAs gave similar patterns, suggesting that this ascertainment bias was largely removed when no close relative lines from those used for discovering InDels were used in diversity analysis. Due to this ascertainment bias, result of our array should be therefore interpreted with caution for diversity analysis.

**Figure 5:**
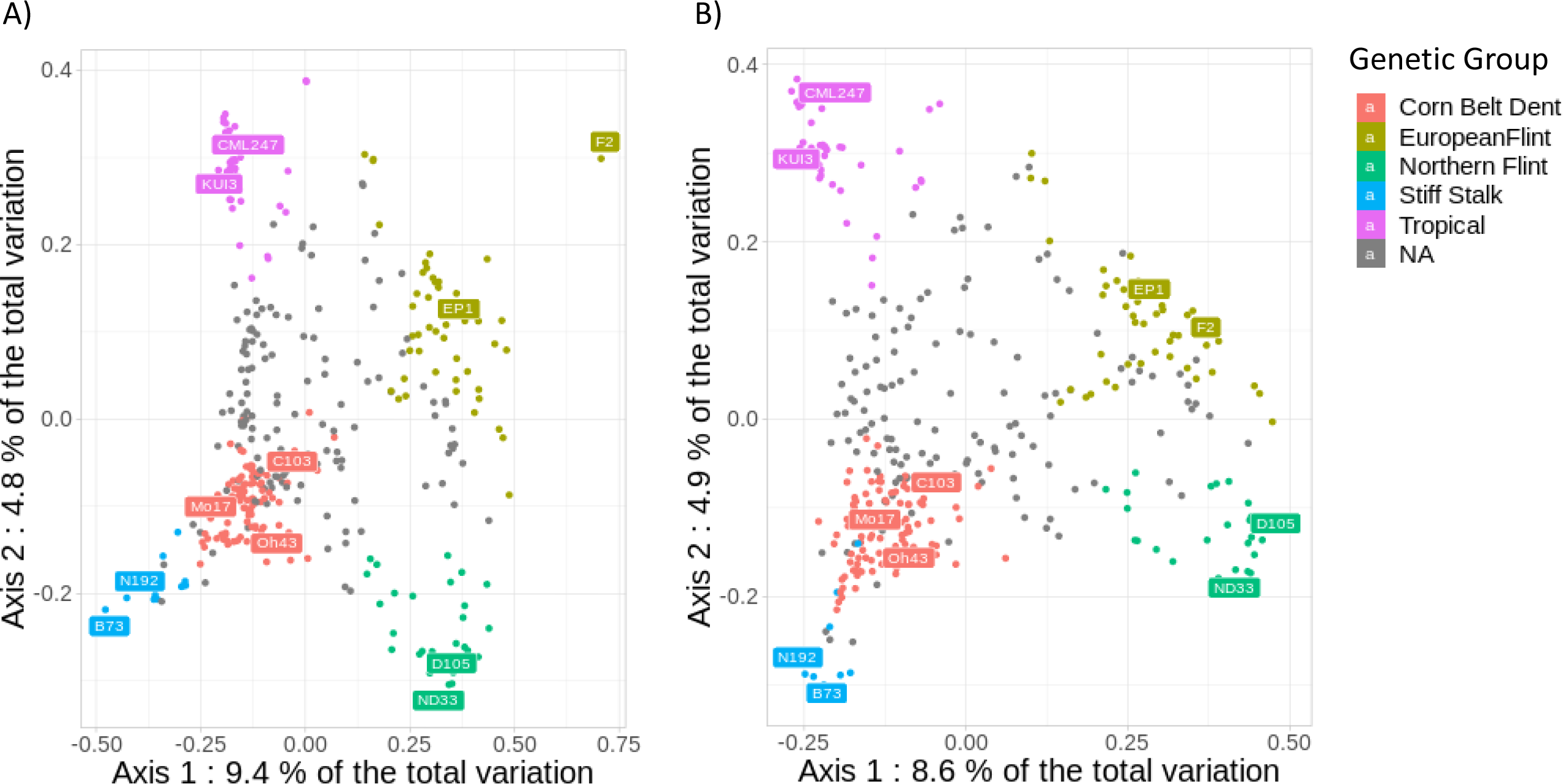
Principal coordinate analysis on the genetic distance between 362 inbred lines from an association panel estimated by A) 57,824 InDels and B) 28,143 SNPs. Colors represent the assignment of the inbred lines to the 5 genetic groups defined by admixture using pre-fixed Panzea SNPs from the 50K Illumina array, when the probability of assignment to a group (membership) was greater than 60%. Inbred lines not assigned to a group were considered admixed and colored gray. The common names of maize accessions, typical of each genetic group, were used.

## Discussion

### 1. An original high throughput approach for genotyping InDels

The comparison of whole genome sequence assemblies is in theory the best approach to identify, precisely and exhaustively, structural variations between two individuals. But even though great progress has been made recently in this area, high-quality, whole genome assembly is still too costly, time-consuming, and computationally intensive to be applied to hundreds of individuals, especially when considering the complexity of the maize genome [20, 58]. Other whole genome sequencing approaches based on alignment of reads on a single reference, and using either “read-depth”, “read-pair”, or “split-read” identification methods [46–50] have mostly been limited to the identification of deletions (i.e. sequences absent from a reference genome). Liu et al., (2015) partially addressed the lack of insertions (i.e. novel sequences compared to a reference genome) by the identification 1,973,746 InDels [4]. Although, among these a majority were very small (85% smaller than 11bp), and the use of PCR markers to genotype them is time-demanding, labor-intensive, and costly at a large-scale level. To avoid this ascertainment bias due to use of a single reference genome to genotype SVs, other studies proposed to call SVs by aligning reads from sequencing on a pan-genome representing the combination of several genomes [14, 20, 22, 52]. However, genotyping InDels with high reliability and call rate by these approaches required at least 30X-50X coverage of the genome to correctly cover their breakpoint and their internal sequence, especially to genotype InDels larger than 50bp [52]. Additionally, aligning reads from a thousand individuals on a pan-genome remained computationally intensive, and therefore required large informatics facilities [52]. In the end, these approaches required to build a pan-genome of high-quality, which remains challenging for a complex genome.

In this paper we describe a new approach combining (i) the ‘accuracy’ of detecting InDels using whole genome assembly, with the detection of 89,150 insertions and 52,175 deletions from the comparison of three newly sequenced and assembled maize inbred line (F2, PH207, and C103) genomes and the public maize B73 AGPv2 reference genome, (ii) and the ‘high-throughput’ genotyping utility provided by SNP arrays. This approach allowed us to genotype, for the first time, thousands of insertion/deletion variants, including PAVs, on a few hundred maize individuals. Genotyping cost per individual using the InDel array was at least 10-20 fold cheaper than any approach based on sequencing for a species with a genome as complex as maize, at a similar level of reliability (> 1000€-2000€ for a 30-50X of a 3Gbp genome *vs* 50€-220€ using Affymetrix^®^ Axiom^®^ array, depending on the number of samples and probes). This genotyping cost did not include bioinformatics analysis. Calling SVs from a pan-genome of a species with a large and complex genome, such as maize, was time-consuming and required bioinformatics skills and large informatics facilities, which are costly and not available in all laboratories. In the contrary, the array could be analyzed rapidly on a laptop using a pipeline already implemented for analyzing SNPs and the Hom2OTV R script developed for analyzing MONO probes. Additionally, the array provided a wet-lab validation of the InDel discovery and allowed the removal of putative genotyping errors from sequencing (particularly for PAVs), due to incomplete or bad genome assembly, as we observed in our study. In the end, the probe content of the InDel array can be largely optimized, either to reduce the size of array (and therefore the cost), or to increase the number of SVs genotyped, without losing reliability (e.g. 200,000 to 300,000 InDels) by filtering out under-performing probes, by strongly reducing the number of probes per InDel (2–3), and by removing false positive InDels. It would also be easy to design an array combining probes targeting InDels and more classical SNPs, outside of InDel sequences.

With the use of breakpoint probes for both insertions and deletions, our approach overcomes some of the limitations of previous CGH or SNP array-based studies, which were only able to call deletions if a few successive probes had lower fluorescent intensity signals [32–35]. Unterseer et al., (2014) genotyped specifically 6,759 small deletions, which were discovered by aligning reads of 30 inbred lines against the B73 genome, but the study did not include any insertions [37]. However, previous CGH and SNP arrays did not design probes to target breakpoints and detected InDels by analyzing the variation of fluorescent intensity signals of ordered probes [32–34]. Consequently, these technologies targeted exclusively low copy regions of the genome, excluding InDels containing repeats, such as transposable elements (TEs) [2, 30, 44]. This is a strong drawback for maize and many other crops since a large part of their sequence is composed of transposable elements [28, 59] which may be highly variable between individuals [4, 24, 60] and may impact phenotypes [61–63]. The use of BP probes allows to target Present/Absent Variations, whose sequence are unique and not present elsewhere in the genome, as well transposable elements, whose internal sequence can be present/absent at one specific locus but also present elsewhere in the genome. Another advantage of genotyping breakpoints is that it provides the ability to genotype the same mutational event across all individuals of the population, as it is highly unlikely that two independent mutational events could lead to the exact same breakpoint. On the contrary, for InDels detected using classical CGH or SNP arrays, it is much harder to identify common InDels among a population of individuals, as we don’t know precisely their breakpoints. Genotyping breakpoints is also very cheap since only one or two probes are needed, which makes the InDel size no longer a limitation for genotyping it accurately, contrary to previous SNP and CGH arrays that rely on fluorescent intensity variation of probes covering the entire InDel sequence [45]. The genotyping of breakpoints by sequencing is possible with a tool like PInDel [50], which has a genotyping mode or BayesTyper [52], but at a much greater cost and with lower call rate compared to the use of a SNP array. Finally, breakpoint probes are codominant markers and allow accurate genotyping of hemizygous individuals (Heterozygous for presence/absence), since their genotyping is based on fluorescent contrast rather than fluorescent intensity variation, which is known to be noisier as with MONO and OTV probes [45].

Although the use of BP probes is clearly the simplest way to genotype InDels using an SNP array, breakpoints are not always available (“no map” approach discovery) or “designable” with 35bp probes, for instance, the cases where sequences of microhomology at breakpoint site were larger than 5bp. In order to genotype the 52,471 InDels without breakpoints and explore the genetic diversity within InDels, we also designed 577,778 internal probes both on monomorphic and polymorphic sites in PARs for both insertions and deletions. To genotype PARs in InDel sequences using SNPs, we took advantage of the already available Affymetrix^®^ algorithms to call Off-Target Variants (OTVs), which can detect variation of fluorescent intensity signals for a single probe (Figure 1C) [36]. This approach was used by [37] who was able to detect 45,974 OTVs on a set of diverse maize inbred lines using a 600K SNP array. Nevertheless, the array was designed in a classical way to target SNPs, and there was no prior evidence that the probes called as OTVs would belong to InDels. Additionally, detecting SNPs in insertions required the assembly of a pan-genome, combining common and specific sequences from different individuals, in order to retrieve SNPs by aligning reads from sequenced lines [14, 20, 22, 52]. In our case, only using OTV probes would have resulted in the elimination of many InDels, since 87,372 of them, including 74,648 insertions, did not have known SNPs within their sequence. In order to avoid this ascertainment bias due to prior knowledge of the presence of SNPs we designed 414,500 MONO probes on putative monomorphic sites within PARs of InDel sequences. This permitted the genotyping of 38,134 supplementary InDels that could not be targeted by OTV or BP probes. This new type of probe required the development of a new algorithm in order to cluster individuals according to their fluorescent intensity variation only, to be able to assign a genotype to each individual (Additional file 2: Figure S7C). A limitation of current workflow is that Affymetrix^®^ algorithms require a larger number of hemizygous individuals to generate high-quality genotype clusters using the OTV and MONO probes. While it was not an issue for maize inbred lines (or individuals from autogamous species) that are mostly homozygous, it was an issue for individuals from allogamous species that are highly heterozygous. By using alternate genotyping techniques or processing a larger number of hemizygous samples, it should be possible to identify hemizygous clusters according to fluorescence intensity from OTV and MONO probes. We observed some clusters that seem incorrectly interpreted as heterozygote for SNPs, although they likely correspond to hemizygous individuals for OTV and MONO probes (Additional file 2: Figure S9B, see below for a more detailed discussion). Alternatively, other algorithms/software based on fluorescent intensity variation of either a single probe or several ordered probes exist and could be used to detect copy number variation for hemizygote individuals [38–43].

In the end, we observed some ascertainment bias using our array (Figure 5). This was due to the fact that our four inbred lines do not well represent the whole genomic diversity of maize, notably missing are tropical lines. As a consequence, it could lead to ascertainment bias by reinforcing the differentiation of inbred lines genetically close to the four inbred lines used to discover InDels [54, 64, 65] as we observed in our diversity analysis for lines close to B73 and F2 (Figure 5 and Figure S13). It could be therefore highly valuable to use more lines for the initial InDel discovery step. Several new individual maize genome assemblies are now available in the public domain and more and more could become available in the future. Our approach could easily be applied to these new genome assemblies to discover new InDels on a larger set of inbred lines representative of maize diversity with the aim to design a new InDel array.

### 2. Reliability of genotyping / calling results

Our approach provides a reliable and reproducible method for genotyping InDels in inbred lines, since (i) the genotypes obtained by array and by sequencing were highly consistent for BP probes (97%) and in a lesser extent with OTV and MONO probes (92% and 88%, respectively), due to the fact that the genome assembly of sequenced lines were incomplete or incorrect, resulting in high error rates for absent calls using GBS; (ii) the average probe genotyping error rate was estimated at 3% (lower for absent calls); (iii) the InDel genotyping errors could be greatly reduced by combining the genotypes of different probes within the InDels (0.02% for 5 probes); (iv) the genotyping results were highly reproducible between DNA replicates of F1 hybrids (95 to 97%, depending on probe type) and between inbred lines (94.8 to 99.4%); and (v) the call rate for individuals was very high (96 to 99%) and can be increased by combining the genotypes of the probes within the InDels (97.7 to 99.9% for 2 and 5 probes, respectively).

Our approach is promising as a method to genotype structural variations in maize, as well as other species with complex genomes. We obtained high metrics, comparable to classical SNP arrays, based on Affymetrix^®^ Axiom^®^ Technologies, even though InDels are more complex to genotype. First, call rates are high and quite similar to those obtained for SNP with the 600K SNP Affymetrix^®^ array (98% against 98.1% in [37]). Nevertheless, we observed a lowest call rate for BP probes (90%). This lowest call rate could be explained by the usage of more relaxing criterion to filter out probes for building array and by the fact that polymorphisms in surrounding sequences of InDel breakpoints have not been taken into account contrary to internal probes. Second, the percentage of BP and OTV probes classified as PHR (94% in both cases) is similar than for 600K SNP Affymetrix^®^ genotyping array (92%) but higher than for 1.2M screening Affymetrix^®^ arrays (~65%) that have been used to select best markers for designing the final 600K SNP Affymetrix^®^ arrays. It is difficult to compare the classification of MONO probes, because the algorithm used (Hom2OTV) is new and quite different from the one used for BP, OTV, and classical SNPs. Third, the reproducibility between DNA replicates of F1 hybrids was high (95 to 97%, depending on probe type), but this is lower than for SNP arrays (~99.5% in [37]). However, the reproducibility was estimated on DNA replicate of F1 hybrids in our study while it was estimated on inbred lines for 600K SNP Affymetrix^®^ array. When we compared genotype of 13 inbred lines originated from different seedlots, reproducibility is close to those of 600K SNP Affymetrix^®^ array (98.3%) and displayed approximatively same reproducibility with 50K SNP Illumina array ([54], Additional file 1: Table S9). This comparison suggested strongly that our lower reproducibility might not be due to genotyping errors but possibly the divergence between the samples for inbred lines and the use of F1 hybrids rather than inbred lines for DNA replicate. Fourth, the Mendelian inheritance between F1 hybrids and their parental lines was lower for our InDel than for SNP array (88% vs 97.6% in [37]) but quite similar considering only homozygous genotypes (95%). This is likely due to the presence of a small number of hemizygous samples since the 16 F1 hybrids were eliminated due to their low call rate (<0.9) and there are only residual hemizygosity for inbred lines. Considering the F1 hybrids for defining BP cluster could improve the delineation of hemizygous cluster and therefore Mendelian inheritance. Note that 600K SNP Affymetrix^®^ in maize was designing by selecting the high confidence probes based on results of a first screening 1.2M SNP Affymetrix^®^ array which could favor reproducibility for this array. Finally, 72% of probes were converted into markers, which is comparable to this 1.2 maize Affymetrix^®^ Axiom^®^ SNP screening arrays (74.9% in [37]). Out of these, 88% were polymorphic for presence/absence. This conversion rate is expected, considering that Affymetrix^®^ Axiom^®^ array analysis pipelines have been optimized for the detection of bi-allelic SNPs and are more sensitive to variations in fluorescent contrast (x-axis) compared to variations in fluorescent intensity (y-axis), which is known to be noisier [36, 45]. Moreover, we did not always follow Affymetrix^®^ recommendations, as we did not filter out probes with a bad design score.

We identified some inconsistencies between genotyping by array (GBA) and genotyping by sequencing (GBS) obtained by aligning probes against our genomes (Table 4). These inconsistencies were higher when GBS called absent for InDels interrogated by OTV and MONO probes (17.1% and 20.2% vs. 4.3% and 5.4%, respectively), although no differences were observed for BP probes (Table 4). These biased inconsistencies towards absence for internal probes seems very high compared to our analysis on the consistencies between probes within Indels. Our analysis of consistencies between probes within InDels showed indeed that genotyping errors produced by the array were close to 3% (Figure 4) and lower for absent calls (Additional file 2: Figure S12). These results suggested that the higher genotyping inconsistencies for GBS absent are due to errors in GBS. GBS errors for absence were well explained by the use of an incomplete genome draft assembly to align probes sequences, and the use of a higher-quality genome could help to reduce these inconsistencies. The probes targeting sequence regions present in one line, but not assembled in their draft genome assemblies, were falsely genotyped absent, but the sample DNA correctly hybridized with the probes, and the InDels were called present with the array. This could also explain why the number of inconsistencies was higher for B73, as all B73 absence genotypes were defined in comparison to draft assemblies. Whereas for the other 3 lines, absence genotypes were defined in comparison with the gold standard B73 genome sequence. The fact that we obtained a better result on OTV probes interrogating InDels discovered in F2 can be explained because we used only SNPs discovered on the B73-F2 pan-genome and not in other genomes. And, the fact that BP probes had similar consistencies for genotyping absent and present calls could be explained by the fact that the BP probes were designed exclusively on B73 reference genome.

We also found that 20,574 InDels were monomorphic and present across all lines, suggesting they represented false positives from regions not assembled in our draft genomes. To reduce this false positive rate, we strongly advise to not only align B73 reads onto each draft genome assembly but to also align reads from each sequenced genome on each other and against itself. This would have several benefits: (i) it would allow to discover even more and higher-quality InDels, as each putative deletion discovered in one sample could potentially benefit from supporting reads from another sample; (ii) this would simplify the identification of InDels common to more than one genotype; and (iii) it would help to identify and eliminate false positive deletions by the alignment of each sample on its own draft assembly.

Nevertheless, the use of incomplete draft genomes does not explain all discrepancies between genotypes obtained by sequencing and by array. First, these discrepancies could also be due to incorrect clustering and assignment of a genotype call (array errors). This was exemplified by some MONO probes classified as SNPs, although the clustering pattern looks like a MONO cluster with a large difference of fluorescence intensity between two clusters (Additional file 2: Figure S9C). A more detailed inspection of the clustering of MONO probes displayed an unexpected cluster pattern (Table 4, Additional file 2: Figure S9D), and OTV probes classified as SNPs (Table 4, Additional file 2: Figure S9A) suggests a wrong assignment of genotypes for the cluster displaying the lowest fluorescent intensity. Similarly, the genome divergence within probe sequences for some inbred lines could result to group those individuals in an OTV cluster, and therefore result in an incorrect absent call. However, these genotyping errors due to bad clustering or genomic divergence between individuals within probes sequences could be strongly reduced by combining genotypes from several probes. As an InDel called by five different probes has a random genotyping error of 5%, we showed by simulation that the genotyping error for that InDel would be reduced to 0.1%, when the most frequent allele among the 5 probes was assigned as genotype of the InDel (Additional file 1: Table S6).

Surprisingly, 4.7% of MONO probes displayed a classical OTV clustering, suggesting that an unknown SNP was targeted by these probes by chance. This high level of polymorphism (1 SNP / 21 bp) was slightly higher than observed by sequencing a small set of diverse lines [66, 67]. It could suggest that PAR genomic regions might have more divergence than other parts of the genome, because these regions were involved in local adaptation by maintaining together favorable combinations of alleles as proposed by [68]. These 15,690 new OTVs are very interesting, since they were discovered by chance on a large set of 445 inbred lines. We could therefore expect that these OTVs have no ascertainment bias, which can be very useful for analyzing genetic diversity within InDels carrying PAR regions. In addition, 20.4% of MONO probes displayed unexpected clustering: one off-target cluster, corresponding to absence of the sequence; one homozygous cluster, corresponding to presence of the sequence; and an unexpected heterozygous cluster (Unexpected MONO 1 in table 4). Considering these “unexpected MONO 1” as true SNPs would indicate a density of 1 SNP every 5 bp, which is not compatible with the level of diversity observed in previous studies of maize [66, 67]. Deeper investigation of these MONO probe clusters identified that for some probes, the unexpected heterozygous cluster is positioned between the presence and absence clusters (Additional file 2: Figure S9B). This suggests that these unexpected heterozygous clusters are identifying inbred samples with only one copy presence (hemizygous genotype). An alternative hypothesis to explain this unexpected pattern is the presence of divergent duplicated sequences, leading to the existence of an artificial heterozygous cluster for SNPs corresponding to the presence of two paralogous sequences. This result suggests therefore that there is probably room to develop genotyping strategies in order to better identify additional clusters corresponding to the presence of hemizygous individuals for both MONO and OTV probes and therefore improve the quality of the genotyping of InDels when using a SNP array.

These potential clustering errors, as well as the incorrect design of some probes, can explain some inconsistent genotypes for presence/absence between probes for the same InDel. Comparison of genotyping across different probes within InDels could help to identify and remove probes displaying highly discordant genotypes, due to errors originating from poor clustering or from poor design. Interestingly, some InDels showed reproducible inconsistent genotypes for presence/absence across their probes in several inbred lines (Additional file 2: Figure S9B). This suggested that this pattern could have a biological origin, with possible rearrangements having occurred several times within the same genomic region in some inbred lines. Following this hypothesis, Gu et al. (2008) observed two different types of rearrangements which could explain our observations [69]: (i) rearrangements with an unique breakpoint in population and therefore common size between individuals resulting to two haplotypes in a population and (ii) rearrangement with non-unique breakpoints, scattered in a genomic region, which resulted in several haplotypes. This hypothesis is also supported in our experiment by the 56% of BP probes classified as OTVs, indicating that FW or/and REV flanking sequence did not hybridize in some lines.

The development of a statistical approach to merge either *a posteriori* the calling results of independent clustering of individual probes or *a priori* the fluorescent intensity signal of successive probes within a InDel could be interesting in order to improve the robustness of InDel genotyping. This would have the advantage to limit the effect of genotyping errors due to poor clustering and to reduce the noise in fluorescent intensity signals. We showed by simulation that assigning the most frequent allele across probes as the genotype reduced genotyping error to 0.7% and 0.1% when 3 and 5 probes were used, respectively. Additionally, it increases the InDel call rate (Additional file 1: Table S6). In the end, it would also help to identify varying haplotypes, representing the complexity of a region in a population. Using multiple probes for calling InDels is therefore highly valuable for improving reliability of InDel genotyping, since it allows putatively to reduce random genotyping error, due to genomic divergence or other causes, removes probes poorly clustered or designed, and identifies more complex rearrangements.

### 3. Conclusions

Our approach, from the sequencing of a few representative genotypes, their genome assembly, the insertion/deletion discovery, and to the design and use of the high-throughput genotyping array was applied to maize as a proof of concept. Our approach allowed us to rapidly create at a reasonable cost a high-throughput SVs genotyping tool for this species. This approach will remain interesting as long as calling large InDels from sequencing, for a large set of individuals, remains un-affordable, bioinformatically challenging, and time-demanding. Nevertheless, our approach could benefit from few improvements based on the knowledge accumulated from this test on maize. First, it could be highly valuable to use more lines for the initial InDel discovery step to avoid ascertainment bias [64] as we observed in our diversity analysis (Figure 5). Using more lines for detecting InDels should also reduce the number of false positives SVs in array due to poor assembly, genotyping error due to genomic divergence between individuals, and help to identify complex rearrangement. Second, even though we did not have any indication that our sequenced data had been contaminated, a contamination cleaning step should be applied to the sequenced data prior to SVs discovery and genome assembly, in order to avoid potential false positive SVs in the final array. Third, aligning reads against the internal sequence of InDels, as well as aligning probes sequences against each genome assemblies, should strongly reduce false positives in the final array. Fourth, improving the pipeline of MONO and OTV probes to call hemizygous genotype from variation of fluorescent data would be very valuable, notably for allogamous species. Fifth, capacity of array could be largely increased to 200,000 or 300,000 InDels without losing reliability by optimizing number of probes per InDels.

To conclude, we developed a “proof of concept” high-throughput and affordable InDel genotyping array, based on the InDels discovered by sequencing on four inbred lines. Our “proof of concept” approach could be easily applied to other species to explore genomic structural variation, notably species with limited sequence data or few genome assemblies available. This could also be interesting for species with greater sequencing resources and where genotyping a large set of individuals is required, such as for
breeding purposes, genome wide association studies, genomic selection, or characterizing SVs in large germplasm. Although our array was not designed to genotype duplications and inversions, our approach could be easily extended to genotype breakpoints of inversions, but further development of the pipeline for genotyping duplications using internal probes would be required. This powerful approach opens the way to studying the contribution of InDels and other SVs to trait variation and heterosis in maize [44] and should contribute to decipher the biological impact of InDels and other SVs at a larger scale in different species.

## Methods

### Sequencing material

Three maize inbred lines, which are key founders of maize breeding programs and originated from three different heterotic groups, had been selected for deep sequencing and InDel discovery: the European Flint line F2 and two American dent lines, PH207 (Iodent) and C103 (Lancaster). For the F2 inbred line, see [20]. For C103 and PH207 inbred lines, DNA was extracted with the NucleoSpin Plant XL, according to the manufacturer’s instructions (Macherey Nagel, Düren, Germany). The DNA concentration was estimated by UV measurement and the quality was checked with an agarose gel electrophoresis. Two library types were sequenced: a 180bp overlapping paired-end library and a 3kb mate-pair library. The paired-end libraries and the sequencing were performed by Integragen (Evry, France) on a HiSeq2000 sequencer (Illumina, San Diego, USA). 412 and 377 million 100bp paired-end reads (33x and 30x) were sequenced respectively for C103 and PH207. The mate-pair libraries were prepared and sequenced at BGI (China) also on HiSeq2000 sequencer (Illumina, San Diego, USA). Raw reads were filtered to remove adaptor sequences, contamination, and low-quality reads. 326 and 316 million 100bp mate-pair reads (26x and 25x) were sequenced, respectively for C103 and PH207. A data set of 473 million B73 inbred line 100bp paired-end reads (35x) with an average insert size of 191bp was downloaded from ftp://ftp.sra.ebi.ac.uk/vol1/fastq/SRR404/SRR404240.

### InDel and PAV discovery

For the deletion discovery step, F2, PH207, and C103 paired-end reads were aligned against B73 AGPv2 genome sequence using novoalign version 3.01.01 (http://www.novocraft.com) (default parameters). Samtools [70] version 0.1.18 was used to coordinate, sort, and retain reads with a mapping quality of at least Q30. Duplicated reads were eliminated using MarkDuplicate from the picardtools suite (http://broadinstitute.github.io/picard) version 1.48. PInDel [50] version 0.2.5a2 was run in parallel on each chromosome to perform multi-genotype calling of deletions. Raw formatted results were converted to VCF (Variant Calling Format) using the script PInDel2vcf. BreakDancer [46] was used in complement PInDel, but only for F2. Deletions shorter than 100bp were discarded. Deletions spanning a B73 assembly gap or located in regions prone to mis-assemblies, such as telomeric, knob, and centromeric regions, were also excluded from further analysis using IntersectBed BEDTools [71] version 2.16.1.

For whole genome sequence reconstruction of F2, PH207, and C103 inbred lines, paired-end and mate-pair reads were used together and assembled using ALLPATHs-LG [72] version R41008 (Additional File 2: Figure S1B). For F2, the script CacheToAllPathsInputs.pl was used to cache the data to use for assembly: 100% of the non-overlapping 230bp insert paired end data set, 100% of the overlapping 170bp insert paired end data set, 30% of the non-overlapping 370bp insert paired end data set, and 100% of the 2.4kb insert mate pair data set. Indeed, only overlapping paired end reads are used by ALLPATHs-LG for building contigs, but the supplementary non-overlapping paired end reads for F2 were used for error correction. RunAllPathsLG was then run for all three genotypes using optional parameters. Details on the sequence library usage during the assembly process are given in Additional file 1: Table S1. For each assembly, the coverage of the gene space was evaluated using BUSCO [73] version 3.0.2 using genome mode and the maize species (-m geno -sp maize).

B73 paired-end reads were successively aligned to ALLPATHs-LG F2, PH207, and C103 genome sequence assemblies (Additional File 2: Figure S1B). The same tools and parameters used to call deletions against the B73 genome were applied to detect B73 deletions against F2, PH207, and C103 genome sequences. These B73 deletions were reciprocally called insertions of F2, PH207, and C103. Only insertions smaller than 100bp were discarded, except those spanning real assembly gaps (with approximate size inferred from paired reads average distance) and not “unsized” gaps like in the B73 genome. When possible, insertions were anchored onto the B73 AGPv2 genome sequence using a dedicated pipeline combining Megablast version 2.2.19 [74] and Age version 0.4 [75]. Again, insertions that could be anchored on the B73 reference and were overlapping regions prone to mis-assemblies such as telomeric, knob, and centromeric regions, were also excluded from further analysis using IntersectBed.

F2 specific sequences coming either from the no map approach (Additional file 2: Figure S1) or from the work of [20] were included as such, without any further filtering.

The multiple references and approaches used during the InDel discovery step led to a set of InDels with various levels of redundancy. Some “intra-tool” redundancy was found (*e.g.* multiple calls found by one tool within the same genotype at highly polymorphic loci). These “ambiguous” calls were systematically identified using the Bedtools suite version 2.16.1 [71] and eliminated. Moreover, for F2 deletions, some “inter-approach” redundancy was also expected and eliminated using intersectBed utility also from the Bedtools suite. When redundancy was found, PInDel calls were preferred to BreakDancer calls, because they had precise breakpoints and contained the calls for PH207 and C103. The same filter was applied to all insertions that could be anchored to the B73 genome sequence. Furthermore, for non-anchored InDels, in order to avoid redundancy in internal genotyping probe design, RepeatMasker (http://www.repeatmasker.org) was used to mask redundant regions by similarity using an iterative approach. First, “ALLPATHs-LG assembly” F2 insertions were masked with “ABySS assembly” F2 insertions (at least 95% of identity) to generate a non-redundant set of F2 insertions. Then C103 insertions were masked with F2 insertions (at 90% of identity), PH207 insertions were masked with C103 and F2 insertions (90%), and finally F2 no map specific sequences were masked with PH207, C103, and F2 insertions (90%).

### Design of Affymetrix® Axiom® array

#### Preparation of sequences for probes for design

To identify presence/absence regions (PARs) within InDel sequences suitable for the design of “off-target” probes, we used the genometools Tallymer utility [76] version 1.5.6 to create two indexes for B73, F2, PH207, and C103: one from their genome assemblies (17-mers with a minimal occurrence of 1) and one from a 5x genome equivalent subset of their raw sequenced data (17-mers with a minimal occurrence of 5). Then B73 genome was iteratively annotated with the script tallymer2gff3.plx (options used: −k 17 −min 35 −occ 1|5 depending on the index) to identify regions not covered by F2, PH207, and C103 kmers. Reciprocally, the two F2 draft genomes, PH207 and C103 ALLPATHs-LG draft genomes were run through the same procedure to identify regions not covered by B73 kmers. The gff files generated by this process were then used in combination with gff files of repeats annotated with RepeatMasker to define PARs of a minimum size of 35bp for each type of InDel and each draft genome.

#### BP preparation

Breakpoints could be targeted by probes (Figure 1A) provided that the nucleotide flanking the breakpoint at the beginning of the deleted sequence was different from the nucleotide right after the end of deleted sequence (and reciprocally on the reverse strand). Type I and type III breakpoints without micro-homology sequence can be submitted for the Affymetrix^®^’ standard design procedure, whereas type II breakpoints have to go through an iterative design process, shifting the sequence by one base on each attempt until reaching a discriminative position. This iterative process stops after 5bp and is also performed by Affymetrix^®^.

#### Probes scoring

All potential probes were evaluated in an *in-silico* analysis to predict their microarray performance. A p-convert value, which arises from a random forest model intended to predict the probability that the SNP will convert on the array, was determined for all probes. The model considers factors including probe sequence, binding energies, and the expected degree of non-specific binding and hybridization to multiple genomic regions. This degree of non-specific binding is estimated calculating 16-mer hit counts, which is the number of times all 16 bp sequences in the 30 bp flanking region from either side of the SNP have a matched sequence in the genome. These scores were generated both for forward and reverse probes. A probeset is recommended if p-convert>=0.6 and there are no expected polymorphisms in the flanking region. A probeset is neutral if p-convert>=0.4, the number of expected polymorphisms in the flanking region is less than 3, and the polymorphisms are further than 21 bp of the variant of interest. Probesets not falling into these two categories are scored as *not recommended*. Probesets that cannot be designed are scored as *not possible*.

#### Probes selection

Concerning OTV and MONO probes, we applied three successive filtering steps. First, we selected only probes classified as recommended and neutral based on their scoring, with no more than one hit on the B73 reference genome for deletion probes, and no hit at all for insertion probes. After this step, 204,213 OTV probes and 18,884,827 MONO probes remained. Secondly, only probes with more than 70% in PARs were kept. An additional filtering step was implemented specifically for MONO probes to optimize probe distribution along the targeted PARs. For this step, PARs were split in 75bp windows using windowmaker (Bedtools) and the MONO probe with the highest p-convert value was selected for each window. If there were InDels with less than 4 MONO probes selected using 75bp windows, these probes were eliminated and a second iteration was attempted, using 50bp windows, followed by a last iteration with 25bp windows. This generated 616,286 probes including BP and OTV probes targeting 108,24 InDels (90% of InDel selected for design). We completed the design by rescuing 6,219 OTV and 53,441 MONO probes from InDels or PARs not targeted by any probes, bringing the total number of probes selected to 675,946 to target 109,292 InDel.

At the last step, duplicated probeset were removed based on their sequence by Affymetrix^®^ during the chip design procedure, leaving 662,772 probeset (105,927 InDels) corresponding to 1,404,570 different probes to be tiled on the array.

### Genotyping of 105k InDels on 480 maize DNA samples

#### Plant material for genotyping

For genotyping, 480 different DNA samples were extracted from leaves following a NaBisulfite method modified from [77, 78]. These 480 samples included 440 inbred lines, 24 highly recombinant inbred lines, and 16 F1 hybrids. Both F1 hybrids (obtained by crossing inbred lines) and their parental inbred lines were genotyped on the array, but seed lots used to produce F1 hybrids and those used to extract DNA for genotyping were different. Among these 480 DNAs, 13 inbred lines were genotyped using two different DNAs from two different seed-lots and were used to evaluate the reproducibility of the genotyping (Additional file 1: Table S9). DNA samples of one F1 hybrid were also genotyped 6 times. Mendelian inheritance was estimated between 12 hybrids F1 derived from 9 different parental lines (Additional file 1: Table S8)

#### Variant calling using Affymetrix® algorithm

Each type of probe had a dedicated algorithm (Additional file 2: Figure S7) to call genotypes, according to expected behavior from the probe design. DNA samples from 480 individuals were hybridized to the array using the Affymetrix^®^ system. The genotyping, sample QC, and marker filtering were performed according to the Axiom^®^ Best Practice genotyping analysis workflow. Genotype calls and classifications were generated from the hybridization signals in the form of CEL files using the Affymetrix^®^ Power Tools (APT) and the SNPolisher package for R, according to the Axiom^®^ Genotyping Solution Data Analysis Guide, and a custom-made R script, Hom2OTV, implemented the algorithm for calling MONO probes.

The APT results were then post-processed using SNPolisher, which is an R package specifically designed by Affymetrix^®^. Marker metrics were generated using the *Ps_Metrics* function. These marker QC metrics were used to classify probesets into 14 categories (Additional file 2: Figure S8) using the *Ps_Classification* and *Ps_Classification_Supplemental* functions, with all default setting for diploid (e.g. HetSO.cut=−0.3, HetvMAF.cut=1.9), except for an empirically determined, more stringent heterozygous variance filter (AB.varY.Z.cut=2.6). Example of clusters from each classification were visualized using the *Ps_Visualization* function (Additional file 2: Figure S8). Variants were preferentially selected as recommended if they were exhibiting stable category assignments with clearly separated clusters. Each variant was ranked into a category (Additional file 2: Figure S8) at each step of the pipeline.

Algorithms used to convert BP and OTV were similar, as BP and OTV probes behaved like classical SNPs. For initial genotype calling, a priori (generic) cluster positions were used, since no information about expected positions was available. A first analysis was performed according to Affymetrix^®^ recommendations. Secondly, the level of inbreeding was taking into account for a posteriori cluster definition, because of the high amount of inbred lines in the panel. This parameter took values from 0 for fully heterozygous to 16 for completely homozygous samples. For OTV and BP algorithms, an inbred penalty of 4 (lower penalty for inbred species) was applied to try to re-labelled probes that fall into categories: CallRateBelowThreshold (CRBT), HomHomResolution (HHR), NoMinorHom (NMH), Other and UnexpectedHeterozygosity, after the first cluster analysis (Additional file 2: Figure S8). Markers that were classified as OTV may also be considered recommended after the *OTV_caller* function has been used to re-label the genotype calls. The SNPolisher *OTV_Caller* function performed post-processing analysis to identify miscalled AB clustering and identify which samples should be in the OTV cluster and which samples should remain in the AA, AB, or BB clusters. Samples in the OTV cluster were re-labelled as OTV. Finally, the recommended markers list is created by combining the list of markers that are classified into the recommended categories (PolyHighResolution (PHR), MonoHighResolution (MHR), and OTV).

BP and OTV probes that exhibited only two clusters (AA or BB and OTV) should fall into the monomorphic classification and be considered as not recommended. A new MONO algorithm was developed (Figure 4), because, unlike traditional SNP genotyping, we only expected two clusters for MONO probes (presence and absence) (Figure 1C). To classify monomorphic sequence genotyping, the *OTV_Caller* function was called, and only MHR and NMH were considered as recommended. Other monomorphic probes are then analyzed with an inbred penalty of 16 (highest level) to re-labelled probes displaying higher-than-expected levels of heterozygosity. Finally, the new function called *Hom2OTV* was implemented to classified probes exhibiting two homozygous clusters, with primarily an intensity difference. This function determined if the intensity difference represents one homozygous cluster (InDel presence) and one OTV cluster (InDel absence), as we expected. There are no parameters in this function. The lower intensity homozygous cluster is recalled as OTV.

#### Evaluation of genotyping quality

We compared the genotyping for 479,027 probes from the InDel array (Genotyping By Array: GBA) with the genotyping from sequencing (Genotyping By Sequencing: GBS) of 4 inbred lines used to discover the InDels: B73, F2, PH207, and C103. Genotyping by sequencing was built from the alignment of probe sequences on the reference genome B73 and the de novo assembly of 3 inbred lines (F2, PH207, and C103) with Blast software. Sequences were considered present in lines when the probes were aligned with less than 5% of mismatch or otherwise considered absent.

Genotyping consistency for B73, F2, PH207, and C103 was calculated between GBS and GBA according to genotype calls “present” or “absent”, produced by GBS (Table 4). For this purpose, Affymetrix^®^ genotyping was converted into these genotypes: present, absent, and hemizygote (1 copy present). Consistency of Presence/Absence genotypes between sequencing and array genotyping was analyzed for four individuals (B73, F2, PH207, C103) according to probe types (BP, OTV, MONO): Number of similar genotypes between GBS and GBA /number of genotype called by GBA and GBS. Note that the seed-lot used for B73 and F2 genotyping is different from the seed-lot used for InDel discovery, but it is the same seed-lot for inbred lines PH207 and C103.

In order to evaluate the consistency of probe genotyping within InDels (Figure 4), we used 362 inbred lines from an association panel representing a wide range of genetic diversity (Camus-Kulandaivelu, 2005; Bouchet et al., 2013). From 479,027 probes, we selected 294,650 polymorphic probes and fully consistent between GBS and GBA in order to limit the genotyping errors due to sequencing. These probes genotyped 72,555 InDels. We then selected 50,648 polymorphic InDels that are genotyped with at least two probes (corresponding to 270,581 probes), and calculated the average frequency of the presence allele across all probes for each InDel and inbred line. For each InDel, we calculated the frequency of inbred lines displaying fully consistent genotypes between probes, *i.e* the proportion of lines where the average frequency across all probes is 0 or 1. We also calculated frequency of inbred lines that have a least one probe with an inconsistent genotype (FreqDiff01), *i.e* the proportion of lines where the average frequency across all probes is not 0 or 1. To evaluate the effect of the probe numbers on the frequency of lines inconsistent within InDels, we analyzed the variation of frequency of lines not fully consistent (FreqDiff01) with relation to the number of probes within the InDels, by estimating median and average FreqDiff01 for each probe count (Figure 4B, Additional file 1: Table S5). To estimate the probe genotyping error rate, we compared this variation to what we could expect for different genotyping error rates (1, 3, 5, and 10%) in 362 lines, genotyped by 10,000 Indels, with the number of probes varying from 2 to 50, using a binomial sampling (Additional file 1: Table S6). For this, we simulated a number of false genotypes among the probes for each InDel and each line using the rbinom function in R, with the following parameters: Number of observation = 362 lines × 10,000 Indels; Number of trials for each observation = Number of probes; Probability of success of each trial = probes genotyping error rate. Using this simulation, we estimated frequency of inconsistent calls among 362,000 simulated genotypes (FreqDiff01) for each probes count, varying from 2 to 50, and compared them with the median and average FreqDiff01 (Figure 4). To evaluate the impact of combining multiple probes for a genotype to correct genotype errors, we used this simulation to estimate the InDel genotyping error rate, if we assign, to an inbred line, the most frequent allele, based on the average allelic frequency of presence (Additional file 1: Table S6). To compare accuracy for genotyping absence and presence using this array, we separated the InDels in four classes, according to their average allelic frequency of absence in 362 inbred lines (0-25%, 25%−50%, 50%−75%, 75%−100%) and compared their median FreqDiff01 (Additional file 2: Figure S12).

To evaluate the reproducibility of the 479,027 probes on the array, we compared the genotypes between 6 DNA replicates from F1 hybrids that originated from crossing B73 and F72. We also compared the genotypes of 13 duplicated inbred lines (A554, A632, A654, B73, C103, CO255, D105, EP1, F2, F252, KUI3, Oh43, and W117) that originated from different seed sources (Additional file 1: Table S9). The genotypes of these 13 duplicated lines were also compared using 43,982 SNPs from the Illumina 50K SNP array.

To evaluate the quality of genotyping, we also analyzed 12 F1 hybrids derived from 9 parental inbred lines Additional file 1: Table S8). We first predicted the genotypes of the 12 F1 hybrids, based on the genotyping of their 2 parental lines, for 46,382 BPs probes, removing OTV calls. These predicted genotypes were then compared with the observed genotypes of the corresponding hybrids: Number of similar genotypes (homozygous or hemizygous) between predicted and observed/Number total of genotypes. BP probes producing missing data or displaying hemizygous genotypes in the parental lies were excluded from the comparison. Note that the seed-lots of the parental inbred lines genotyped may have been different from the seed-lots used for producing the F1 hybrids.

### Diversity analysis

We performed diversity analysis on 362 inbred lines from an association panel representing a wide range of diversity [55, 57], obtained using InDels genotyped on our InDels Affymetrix^®^ Axiom^®^ array and using SNPs genotyped from the Illumina 50K SNP array [54]. The genotypes of InDels were treated as bi-allelic “present” and “absent”.

To perform diversity analysis, we first selected 237,629 probes among the 479,027 probes for which (i) the clustering observed were consistent with expectation (Table 3) and (ii) for which genotypes produced by our array for the 4 lines used for discovering the InDels were fully consistent with the genotyping, based on the alignment of the probes on the genome assemblies using BLAST software. We filtered out 219,068 probes based on their genotyping quality (missing data rate below 20%, heterozygous rate below 15% and minor allele frequency above 5%). In the end, we selected a single, best probe for each InDel, leading to a set of 57,824 probes genotyping 57,824 InDels to analyze diversity in 362 inbred lines.

We estimated two kinship matrices between 362 lines using “identity by descent” estimators (IBD) based on 57,824 InDels and on 28,143 prefixed Panzea SNPs from the Illumina 50K (Figure 5). Kinship matrices were estimated with the “ibd” function in the R package GenABEL [79]. We performed correlation between IBD values estimated with SNP and InDel polymorphisms. Genetic structuration was estimated using only the 28,143 Panzea SNPs with admixture software [80]. We selected the admixture results for five genetic groups (Q=5), since it corresponded to the number of genetics groups defined in previous studies using the Panzea SNPs from the Illumina 50K [55]. Lines were assigned to one genetic group, given that the probability of assignment to the groups was greater than 0.6, whereas lines below this threshold were considered “admixed”. In order to compare genetic structuration based on InDels and SNPs, we performed Principal Coordinate Analysis (PcoA) on genetic distance between lines with (362 lines) and without F2 and B73 (360 lines) based on their dissimilarity (1-IBD) using InDels. Each line was plotted on the first two planes of PcoA and colored according to the assignment to the 5 genetics groups (Figure 5).

## List of abbreviations

GBA: Genotyping by array
GBS: Genotyping by sequencing
SNP: Single nucleotide polymorphism
InDel: Insertion / Deletion
BP: Breakpoint
MONO: Monomorphic
OTV: Off Target Variant
QC: Quality control
PHR: Poly High Resolyion
VCF: Variant Call Format
PAR: Presence / Absence Region
PAV: Presence / Absence Variant
SV: Structural variant
CNV: Copy Number Variant
TE: Transposable Element
CGH: Comparative Genomic Hybridization
NGS: Next Generation Sequencing
FW: Forward
REV: Reverse
NAM: Nested Association Mapping
DNA: Deoxyribonucleic Acid
PCR: Polymerase Chain Reaction
PcoA: Principal Coordinate Analysis
Mbp: Millions of Base Pairs
bp: base pair
FreqDiff01: Frequency of lines not fully consistent between probes within InDel

## Declarations

## Ethics approval and consent to participate

Not applicable

## Consent to publish

Not applicable

## Availability of data and materials

The array content is available at https://doi.org/10.15454/DWB4UT

## Competing interests

Ali Pirani is an employee of Thermo Fisher Scientific (formerly Affymetrix®).

## Funding

This work was supported by the project CNV-MAIZE (ANR-10-GENM-003) and the project Investement for the future AMAIZING ANR-10-BTBR-01 (ANR-PIA AMAIZING) and France Agrimer. PhD student C. Mabire is jointly funded by the program CNV4sel in the framework of metaprogram Selgen and by the Plant Biology and Breeding department of the French National Institute for Agricultural Research (INRA).

## Authors’ contributions

SDN designed and supervised the study and conducted CNVMaize project. CM, JD, and SDN drafted and corrected the manuscript, CV and JJ corrected the manuscript. NR, SDN, JPP, and SP conceived the array, AP, SDN, JJ and JD designed the array. AP developed the Affymetrix® genotype calling software and preformed the InDel genotype calling. JPP, JJ, and CV contributed to the sequencing; JD, AD, HR, and JJ performed the InDel discovery, JD and JJ build genome assemblies, JJ and AD discovered SNP within the InDels. CM evaluated the quality of genotyping and conducted the genetic diversity analysis.DM and VC did DNA extraction and prepared the samples for arrays genotyping. All authors have read and approved the manuscript.

## Acknowledgements

We are very grateful to Patrick Schnable and Cheng-Ting “Eddy” Yeh for providing a subset of Presence/Absent Variants coming from their RNAseq and Sequence capture approach and for his helpful discussion. We are very grateful to Alain Charcosset for their contribution to the choice of inbred lines genotyped by the array and for his helpful discussion and comments on the manuscript.

## Additional Files

### Additional file 1

**Table S1:** Summary of sequencing data used during the assembly process provided by ALLPATHS-LG; **Table S2:** Classification by the Affymetrix^®^ pipeline of 84,994 BP probes based on cluster number, separation, variance, and call rate. A) Probes recommended for genotyping, B) Probes not recommended for genotyping; **Table S3**: Classification by the Affymetrix^®^ pipeline of 163,278 OTV probes based on cluster number, separation, variance, and call rate. A) Probes recommended for genotyping B) Probes not recommended probes for genotyping; **Table S4:** Classification by the Affymetrix^®^ pipeline of 414,500 MONO probes, based on cluster number, separation, variance, and call rate. A) Probes recommended for genotyping B) Probes not recommended for genotyping; **Table S5:** Effect of probe number within InDels on average percentage of missing data, of genotypes absent and genotypes not fully concordant; **Table S6:** Simulation of genotyping error rates for 362 lines and 10,000 InDels called by various numbers of probes with a probe genotyping error rate ranging from 1% to 10%; **Table S7:** Comparison of reproducibility between 5 DNA replicates of hybrid F1 according to probes type and observed clustering; **Table S8:** Mendelian inheritance of 12 hybrids F1 derived from 9 different parental inbred lines for 46,382 BP probes passing Affymetrix quality control and polymorphic; **Table S9**: Comparison of the reproducibility of InDels and SNP genotyping between 13 maize varieties replicated on 50K Illumina SNP and Affymetrix^®^ Axiom^®^ InDel arrays

### Additional file 2

**Figure S1:** Description of two approaches used to discover InDels using resequencing data of DNA; **Figure S2:** Number and complementarity of A) deletions and B) insertions regarding B73 reference genome discovered between F2, PH207 and C103 inbred lines and B73; **Figure S3:** Schematic representation of four different breakpoint types identified by PINDEL at InDel breakpoints according to the presence of micro-homology sequence or not in place of the deleted sequence; **Figure S4:** Distribution of probe number per InDel for 105,927 InDels genotyped with the array; **Figure S5:** Relationship between probes number genotyping the InDel and A) the InDel length B) cumulated length of specific sequence (PARs) within InDel; **Figure S6:** Variation of probes and InDels density across the 10 maize chromosomes; **Figure S7:** Three dedicated Affymetrix pipelines used for calling InDel polymorphisms from the fluorescent intensity variation of BP probes (A), OTV probes (B) and MONO probes (C); **Figure S8:** Example of clustering based on probes fluorescence (intensity in y-axis and contrast in x-axis), for 14 different classifications of probes assigned by the Affymetrix^®^ algorithm; **Figure S9:** Example of clustering for 6 randomly probes in different classifications; **Figure S10:** Variation of the distribution of the average consistency rate (%) of InDels between expected and observed genotyping of probes according to number of probes within the InDel; **Figure S11:** Haplotype of two InDels genotyped with multiple probes (in column) for 362 individuals (in rows); **Figure S12:** Effect of average frequency of absence across 362 lines on consistencies between probes genotyping within InDels; **Figure S13:** Comparison of kinship between 362 inbred lines estimated with 57,824 InDels and with 28,143 SNPs from the 50K Illumina genotyping array; **Figure S14:** Principal coordinate analysis on the genetic distance between 360 inbred lines from an association panel (B73 and F2 were excluded) estimated by A) 57,824 InDels and B) 28,143 SNPs.

## Notes

#### Summary of Updates

In this new version, we added an estimation of genotyping error rate and several metrics on array (reproducibility, mendelian inheritance...). We added a discussion about the interest of this array compared to sequencing.

https://doi.org/10.15454/DWB4UT

