## Additional file 1: Supplementary Table for "High throughput genotyping of structural variations in a complex plant genome using an original Affymetrix® Axiom® array"

### ADDITIONAL FILE 1 : SUPPLEMENTARY TABLES

*Table S1: Summary of sequencing data used during the assembly process provided by ALLPATHS-LG.*

| Inbred lines | Type | Distance R1-R2 | n reads | % used | scov | n pairs | pcov |
|---|---|---|---|---|---|---|---|
| F2 | PE | -36 (+/- 27) | 516,844,994 | 18.2 | 8.5 | 174,374,819 | 15.7 |
| | PE | 26 (+/- 38) | 972,86,460 | 32.9 | 5.7 | 27,367,318 | 11.8 |
| | PE | 181 (+/- 55) | 219,355,916 | 22.3 | 8.6 | 47,582,696 | 33.3 |
| | | Total PE | 833,487,370 | 21 | 31.3 | 174,374,819 | 76.3 |
| | MP | 2153 (+/- 262) | 565,822,128 | 17.2 | 16.4 | 18,850,205 | 108.4 |
| C103 | PE | -37 (+/- 15) | 823,506,904 | 31.9 | 36.4 | 246,941,948 | 61.9 |
| | MP | 2898 (+/- 150) | 652,864,236 | 9.6 | 8.6 | 11,167,486 | 69.7 |
| PH207 | PE | -26 (+/- 16) | 754,615,040 | 25.5 | 27.9 | 217,832,024 | 59.1 |
| | MP | 3053 (+/- 177) | 631,282,744 | 8.5 | 7.3 | 10,592,560 | 69 |

*Type: Type of library with PE=Paired End and MP = Mate Paired ; Distance R1-R2 = Average Insert Size with standard deviation in bracket ;n reads (number of reads in input) %_used (% of reads assembled) ; scov (sequence coverage) ; n_pairs (number of valid pairs assembled) ; pcov (physical coverage Sequence).*

*Table S2: Classification by the Affymetrix® pipeline of 84,994 BP probes based on cluster number, separation, variance, and call rate. A) Probes recommended for genotyping, B) Probes not recommended for genotyping.*

**A) Recommended probes**

| Affymetrix Classification | Pipeline Step † | Number of probes | Percentage of probes | Number of InDels |
|---|---|---|---|---|
| AAvarianceX | AxiomGT1 | 123 | 0.27% | 123 |
| AAvarianceX | Inbred | 139 | 0.30% | 139 |
| AAvarianceX | OTV | 242 | 0.52% | 241 |
| AAvarianceY | OTV | 760 | 1.64% | 752 |
| ABvarianceX | AxiomGT1 | 36 | 0.08% | 36 |
| ABvarianceX | Inbred | 2 | 0.00% | 2 |
| ABvarianceX | OTV | 25 | 0.05% | 25 |
| ABvarianceY | OTV | 223 | 0.48% | 221 |
| BBvarianceX | AxiomGT1 | 122 | 0.26% | 122 |
| BBvarianceX | Inbred | 122 | 0.26% | 122 |
| BBvarianceX | OTV | 254 | 0.55% | 253 |
| BBvarianceY | OTV | 759 | 1.64% | 749 |
| PolyHighResolution | AxiomGT1 | 14,932 | 32.19% | 12,981 |
| PolyHighResolution | Inbred | 4,653 | 10.03% | 4,484 |
| PolyHighResolution | OTV | 23,990 | 51.72% | 20,061 |
| | **Total** | 46,382 | 100% | 34,232 |

**B) Not Recommended probes**

| Affymetrix Classification | Pipeline Step† | Number of probes | Percentage of probes | Number of InDels |
|---|---|---|---|---|
| CallRateBelowThreshold | inbred | 2,915 | 7.55% | 2,845 |
| HomHomResolution | inbred | 2,228 | 5.77% | 2,185 |
| MonoHighResolution | inbred | 4,972 | 12.88% | 4,676 |
| NoMinorHom | inbred | 2,583 | 6.69% | 2,457 |
| Other | inbred | 20,533 | 53.18% | 18,302 |
| UnexpectedHeterozygosity | inbred | 21 | 0.05% | 21 |
| CallRateBelowThreshold | OTV | 4,345 | 11.25% | 4,256 |
| HomHomResolution | OTV | 2 | 0.01% | 2 |
| MonoHighResolution | OTV | 112 | 0.29% | 111 |
| NoMinorHom | OTV | 304 | 0.79% | 300 |
| Other | OTV | 378 | 0.98% | 373 |
| OTV | OTV | 211 | 0.55% | 210 |
| UnexpectedHeterozygosity | OTV | 8 | 0.02% | 8 |
| | **Total** | 38,612 | 100% | 31,049 |

*†Pipeline step corresponds to the step at which BP probes were classified as described in figure S7: AxiomGT1, Inbred, OTV for steps "Axiom GT1","AxiomGT1 inbred penalty 4", "OTV caller" in figure S7, respectively.*

*Table S3: Classification by the Affymetrix® pipeline of 163,278 OTV probes based on cluster number, separation, variance, and call rate. A) Probes recommended for genotyping B) Probes not recommended probes for genotyping.*

**A) Recommended probes**

| Affymetrix Classification | Pipeline Step[ǂ] | Number of probes | Percentage of probes | Number of InDels |
|---|---|---|---|---|
| AAvarianceX | AxiomGT1 | 102 | 0.11% | 102 |
| AAvarianceX | OTV | 784 | 0.81% | 666 |
| AAvarianceY | OTV | 1,291 | 1.33% | 1,100 |
| ABvarianceX | AxiomGT1 | 20 | 0.02% | 19 |
| ABvarianceX | Inbred | 1 | 0.00% | 1 |
| ABvarianceX | OTV | 145 | 0.15% | 139 |
| ABvarianceY | OTV | 658 | 0.68% | 570 |
| BBvarianceX | AxiomGT1 | 127 | 0.13% | 124 |
| BBvarianceX | OTV | 1,030 | 1.06% | 903 |
| BBvarianceY | OTV | 1,352 | 1.40% | 1,128 |
| HomHomResolution | OTV | 71 | 0.07% | 71 |
| MonoHighResolution | OTV | 506 | 0.52% | 467 |
| PolyHighResolution | AxiomGT1 | 11,485 | 11.86% | 5,179 |
| PolyHighResolution | Inbred | 4913 | 5.07% | 3,285 |
| PolyHighResolution | OTV | 74,382 | 76.79% | 13,097 |
| | **Total** | 96,867 | 100% | 15,064 |

**B) Not recommended probes**

| Affymetrix Classification | Pipeline Step[ǂ] | Number of probes | Percentage of probes | Number of Indels |
|---|---|---|---|---|
| NoMinorHom | inbred | 5,746 | 8.65% | 3,705 |
| UnexpectedHeterozygosity | inbred | 32 | 0.05% | 31 |
| AAvarianceY | OTV | 1 | 0.00% | 1 |
| CallRateBelowThreshold | OTV | 31,066 | 46.78% | 10,471 |
| MonoHighResolution | OTV | 522 | 0.79% | 486 |
| NoMinorHom | OTV | 2,758 | 4.15% | 2,218 |
| Other | OTV | 25,086 | 37.77% | 9,297 |
| OTV | OTV | 686 | 1.03% | 637 |
| PolyHighResolution | OTV | 2 | 0.00% | 2 |
| UnexpectedHeterozygosity | OTV | 512 | 0.77% | 470 |
| | **Total** | 66,411 | 100% | 14,293 |

*[ǂ]Pipeline step corresponds to the step at which OTV probes were classified as described in figure S7. AxiomGT1, Inbred, and OTV for steps "Axiom GT1", "AxiomGT1 inbred penalty 4"; OTV: third step "OTV caller" in figure 7, respectively*

*Table S4: Classification by the Affymetrix® pipeline of 414,500 MONO probes, based on cluster number, separation, variance, and call rate. A) Probes recommended for genotyping B) Probes not recommended for genotyping.*

**A) Recommended probes**

| **Affymetrix Classification**[*] | **Pipeline Step** | **Number of probes** | **Percentage of probes** | **Number of InDels** |
|---|---|---|---|---|
| AAvarianceX | Inbred | 11 | 0.00% | 11 |
| AAvarianceX | Poly | 309 | 0.09% | 305 |
| AAvarianceY | Poly | 384 | 0.11% | 382 |
| ABvarianceX | Poly | 84 | 0.03% | 84 |
| ABvarianceY | Poly | 256 | 0.08% | 253 |
| BBvarianceX | Inbred | 3 | 0.00% | 3 |
| BBvarianceX | Poly | 156 | 0.05% | 156 |
| BBvarianceY | Poly | 878 | 0.26% | 856 |
| HomHomResolution | Inbred | 3,867 | 1.15% | 3622 |
| HomHomResolution | Poly | 4 | 0.00% | 4 |
| MonoHighResolution | Mono | 19,771 | 5.89% | 13300 |
| MonoHighResolution | Poly | 27,165 | 8.09% | 14222 |
| NoMinorHom | Mono | 25,820 | 7.69% | 16716 |
| Other | Inbred | 213,759 | 63.66% | 55225 |
| OTV | Inbred | 11,317 | 3.37% | 8933 |
| PolyHighResolution | Inbred | 16,394 | 4.88% | 11036 |
| PolyHighResolution | Poly | 15,600 | 4.65% | 12531 |
| | **Total** | 335,778 | 100% | 63597 |

**B) Not recommended probes**

| Affymetrix Classification* | Pipeline Step† | Number of probes | Percentage of probes | Number of InDels |
|---|---|---|---|---|
| CallRateBelowThreshold | Poly | 7,401 | 9.40% | 6466 |
| NoMinorHom | Poly | 3,232 | 4.11% | 2905 |
| Other | Poly | 17,850 | 22.67% | 13539 |
| OTV | Poly | 3,194 | 4.06% | 2912 |
| UnexpectedHeterozygosity | Poly | 23 | 0.03% | 23 |
| AAvarianceX | Inbred | 106 | 0.13% | 106 |
| AAvarianceY | Inbred | 144 | 0.18% | 143 |
| ABvarianceX | Inbred | 1 | 0.00% | 1 |
| ABvarianceY | Inbred | 18 | 0.02% | 18 |
| BBvarianceX | Inbred | 32 | 0.04% | 32 |
| BBvarianceY | Inbred | 138 | 0.18% | 138 |
| CallRateBelowThreshold | Inbred | 2,717 | 3.45% | 2583 |
| HomHomResolution | Inbred | 5,380 | 6.83% | 4940 |
| MonoHighResolution | Inbred | 22,627 | 28.74% | 15,509 |
| NoMinorHom | Inbred | 3,404 | 4.32% | 3,160 |
| Other | Inbred | 8,095 | 10.28% | 7,126 |
| OTV | Inbred | 786 | 1.00% | 776 |
| PolyHighResolution | Inbred | 3,574 | 4.54% | 3,328 |
| | **Total** | 78,722 | 100% | 38,307 |

† *Pipeline step corresponds to the step at which MONO probes were classified as described in figure S7: Poly, Mono and Inbred for step "Polyploid analysis OTV caller", "Monomorphic analysis OTV caller" and "AxiomGT1 inbred penalty 16", respectively*

**Note that criteria for classifying probes according to the quality of their clustering was not defined for the Hom2OTV software. For MONO probes called by the Hom2OTV algorithm, we displayed Affymetrix classification from the step Inbred.*

*Table  S5: Effect of probe number within InDels on average percentage of missing data, of genotypes absent and genotypes not fully concordant.*

| Number of probes | Average FreqDiff01 (percent) | Median FreqDiff01 (percent) | Number of InDels | Percentage of genotypes absent among all genotypes | Average missing data rate |
|---|---|---|---|---|---|
| 2 | 9.9% | 1.2% | 11164 | 57% | 2.35% |
| 3 | 18.7% | 6.6% | 7297 | 50% | 0.09% |
| 4 | 22.4% | 12.4% | 8014 | 50% | 0.03% |
| 5 | 25.1% | 16.3% | 5783 | 49% | 0.02% |
| 6 | 28.0% | 21.0% | 3923 | 49% | 0.01% |
| 7 | 29.5% | 22.1% | 2576 | 46% | 0.02% |
| 8 | 31.6% | 26.0% | 1863 | 46% | 0.03% |
| 9 | 33.5% | 28.7% | 1402 | 45% | 0.00% |
| 10 | 35.9% | 32.0% | 1145 | 45% | 0.00% |
| 11 | 38.3% | 37.8% | 931 | 44% | 0.00% |
| 12 | 38.3% | 35.9% | 725 | 45% | 0.00% |
| 13 | 39.3% | 37.0% | 605 | 44% | 0.00% |
| 14 | 41.5% | 38.7% | 503 | 45% | 0.00% |
| 15 | 44.7% | 45.0% | 450 | 43% | 0.00% |
| 16 | 44.7% | 46.1% | 339 | 43% | 0.00% |
| 17 | 44.0% | 43.6% | 359 | 44% | 0.00% |
| 18 | 43.7% | 42.8% | 271 | 43% | 0.00% |
| 19 | 47.2% | 46.7% | 238 | 43% | 0.00% |
| 20 | 48.0% | 48.5% | 242 | 45% | 0.00% |

*Note that average missing data were estimated on real data after combining genotyping of all probes within InDel.*

*Table S6: Simulation of genotyping error rates for 362 lines and 10,000 InDels called by various numbers of probes with a probe genotyping error rate ranging from 1% to 10%.*

| Number of probes | Genotyping Error rate+ | | Percentage of genotype unassigned* | Percentage of genotype not totally consistent across probes within an InDel |
|---|---|---|---|---|
| | Probe | InDel | | |
| 1 | 1% | 0.997% | 0.00% | 1.0% |
| 2 | | 0.009% | 1.98% | 2.0% |
| 3 | | 0.029% | 0.00% | 3.0% |
| 4 | | 0.000% | 0.06% | 3.9% |
| 5 | | 0.001% | 0.00% | 4.9% |
| 6 | | 0.000% | 0.00% | 5.9% |
| 7 | | 0.000% | 0.00% | 6.8% |
| 8 | | 0.000% | 0.00% | 7.7% |
| 9 | | 0.000% | 0.00% | 8.6% |
| 10 | | 0.000% | 0.00% | 9.6% |
| 1 | 3% | 2.999% | 0.00% | 3.0% |
| 2 | | 0.088% | 5.82% | 5.9% |
| 3 | | 0.268% | 0.00% | 8.8% |
| 4 | | 0.011% | 0.51% | 11.4% |
| 5 | | 0.026% | 0.00% | 14.1% |
| 6 | | 0.001% | 0.05% | 16.7% |
| 7 | | 0.002% | 0.00% | 19.2% |
| 8 | | 0.000% | 0.01% | 21.6% |
| 9 | | 0.000% | 0.00% | 23.9% |
| 10 | | 0.000% | 0.00% | 26.2% |
| 1 | 5% | 5.004% | 0.00% | 5.0% |
| 2 | | 0.250% | 9.49% | 9.7% |
| 3 | | 0.724% | 0.00% | 14.3% |
| 4 | | 0.047% | 1.36% | 18.6% |
| 5 | | 0.116% | 0.00% | 22.6% |
| 6 | | 0.009% | 0.21% | 26.5% |
| 7 | | 0.020% | 0.00% | 30.2% |
| 8 | | 0.001% | 0.03% | 33.6% |
| 9 | | 0.003% | 0.00% | 37.0% |
| 10 | | 0.000% | 0.01% | 40.2% |
| 1 | 10% | 9.991% | 0.00% | 10.0% |
| 2 | | 0.991% | 18.01% | 19.0% |
| 3 | | 2.794% | 0.00% | 27.1% |
| 4 | | 0.368% | 4.88% | 34.4% |
| 5 | | 0.855% | 0.00% | 41.0% |
| 6 | | 0.128% | 1.47% | 46.8% |
| 7 | | 0.273% | 0.00% | 52.2% |
| 8 | | 0.041% | 0.46% | 57.0% |
| 9 | | 0.089% | 0.00% | 61.2% |
| 10 | | 0.014% | 0.15% | 65.1% |

**Note that genotype remain unassigned when the frequency of two alleles were equal. +Genotypes of each InDel were obtained by assigning the most frequent allele across the different probes for each line of panel.*

*Table S7: Comparison of reproducibility between 5 DNA replicates of hybrid F1 according to probes type and observed clustering.*

| | | BP | MONO | OTV | BP | MONO | OTV |
|---|---|---|---|---|---|---|---|
| | | Number of probes | | | Median consistencies rate† | | |
| Observed clustering* | BP | 20,370 | - | - | 0.955 | - | - |
| | MONO | - | 212,434 | 502 | - | 0.968 | 0.983 |
| | monomorphic | - | 27,586 | 4 | - | 0.999 | 1.000 |
| | OTV | 26,012 | 15,690 | 78,799 | 0.940 | 0.965 | 0.962 |
| | OTV_Type2 | - | 68,562 | - | - | 0.978 | - |
| | SNP | - | 1,981 | 17,562 | - | 0.966 | 0.946 |
| | SNP_Type2 | - | 9,525 | - | - | 0.992 | - |

*Observed clustering corresponds to different clustering classes for each probe types identified in table 3.*

† *Median consistency rate was estimated using pairwise comparison of genotype between 5 DNA replicates.*

*Table S8: Mendelian inheritance of 12 hybrids F1 derived from 9 different parental inbred lines for 46,382 BP probes passing Affymetrix quality control and polymorphic.*

| Hybrids F1 | Proportion of genotypes observed in F1 hybrids different from those predicted from their two parental inbred lines according to genotypes in hybrids* | | | | Proportion of missing loci* |
|---|---|---|---|---|---|
| | Hemizygous | Homozygous | Homozygous "Absent" | All | |
| F72 x B73 | 0,22 | 0,05 | 0,02 | 0,11 | 0,14 |
| B73 x W117 | 0,24 | 0,12 | 0,20 | 0,15 | 0,13 |
| MO17 x B73 | 0,24 | 0,04 | 0,02 | 0,12 | 0,11 |
| PH207 x B73 | 0,22 | 0,06 | 0,02 | 0,12 | 0,11 |
| B73 x D105 | 0,23 | 0,04 | 0,04 | 0,11 | 0,14 |
| B73 x EP1 | 0,22 | 0,04 | 0,03 | 0,11 | 0,14 |
| F252 x F2 | 0,25 | 0,05 | 0,01 | 0,11 | 0,17 |
| MO17 x F2 | 0,25 | 0,05 | 0,02 | 0,12 | 0,16 |
| PH207 x F2 | 0,25 | 0,09 | 0,01 | 0,14 | 0,17 |
| W117 x F2 | 0,25 | 0,08 | 0,02 | 0,12 | 0,17 |
| F2 x D105 | 0,25 | 0,05 | 0,01 | 0,10 | 0,17 |
| F2 x EP1 | 0,23 | 0,04 | 0,01 | 0,08 | 0,16 |
| Average | 0,24 | 0,06 | 0,03 | 0,12 | 0,15 |
| Median | 0,24 | 0,05 | 0,02 | 0,12 | 0,15 |

* *Missing loci: Loci displaying missing data, hemizygous, or off-target genotypes for one of the parental line in addition to loci displaying missing data or off-target genotypes in hybrids F1 were not considered for comparison.*

† *Genotype called "absent" in hybrid F1*

*Table S9: Comparison of the reproducibility of InDels and SNP genotyping between 13 maize varieties replicated on 50K Illumina SNP and Affymetrix® Axiom® InDel arrays.*

| | **% of difference in genotyping*** | |
|---|---|---|
| **Variety** | **50K SNP array** | **InDel array** |
| A554 | 4.4% | 4.5% |
| A632 | 1.7% | 2.1% |
| A654 | 1.6% | 1.4% |
| B73 | 0.0% | 5.2% |
| C103 | 0.2% | 0.6% |
| CO255 | 1.5% | 0.9% |
| D105 | 1.7% | 0.7% |
| EP1 | 1.7% | 2.0% |
| F2 | 1.6% | 1.7% |
| F252 | 3.1% | 0.9% |
| KUI3 | 6.0% | 5.2% |
| Oh43 | 0.3% | 2.6% |
| W117 | 1.6% | 0.8% |

**Percentage of genotypes different between replicates from a same variety. Note that DNA samples from replicated varieties originated from different seed sources*
