## Additional file 2: Supplementary figures for "High throughput genotyping of structural variations in a complex plant genome using an original Affymetrix® Axiom® array"

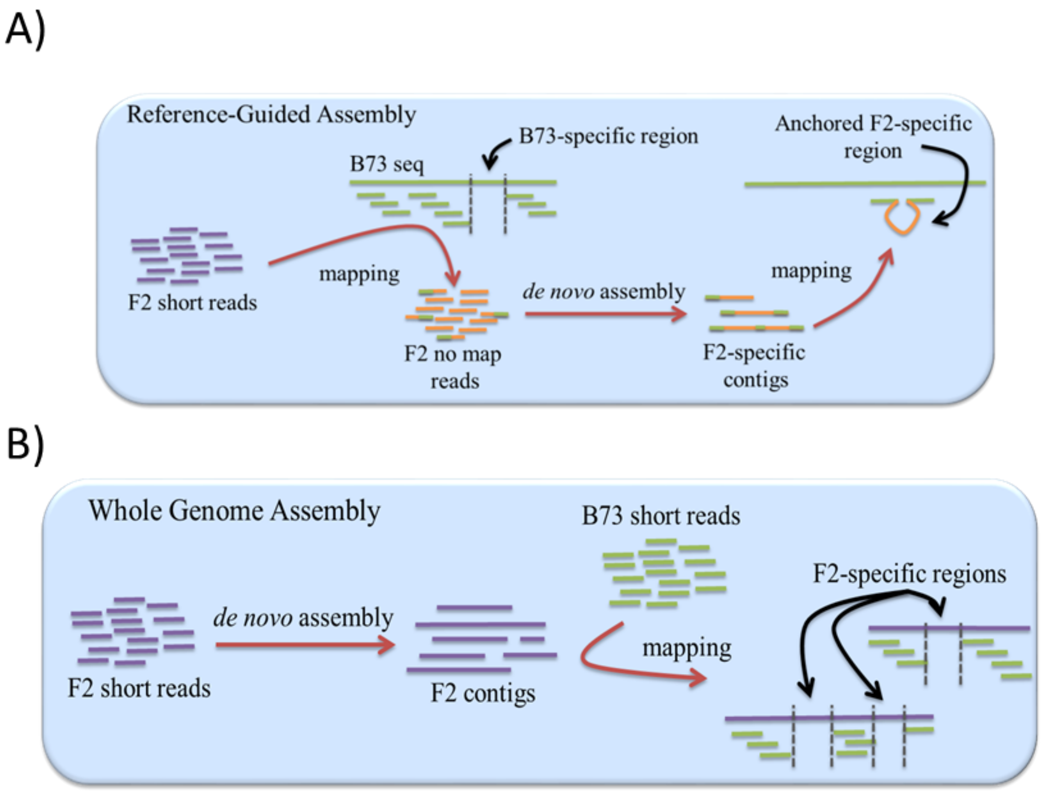

*Figure S1: Description of two approaches used to discover InDels using resequencing data of DNA.*

A) reference guided assembly ("no map" approach) used only on F2 and B) whole genome assembly used on F2, PH207 and C103.

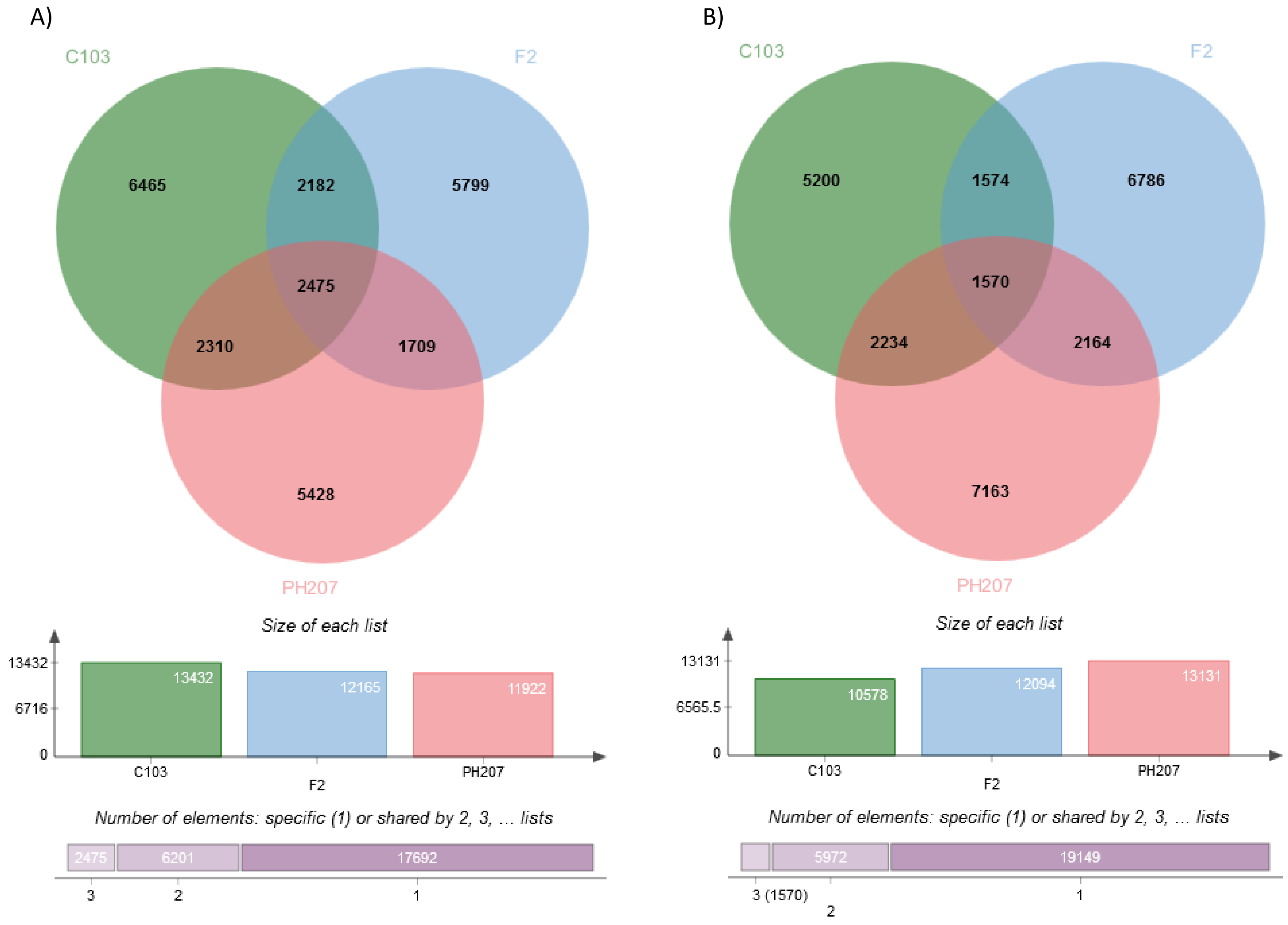

*Figure S2: Number and complementarity of A) deletions and B) insertions regarding B73 reference genome discovered between F2, PH207 and C103 inbred lines and B73*

34

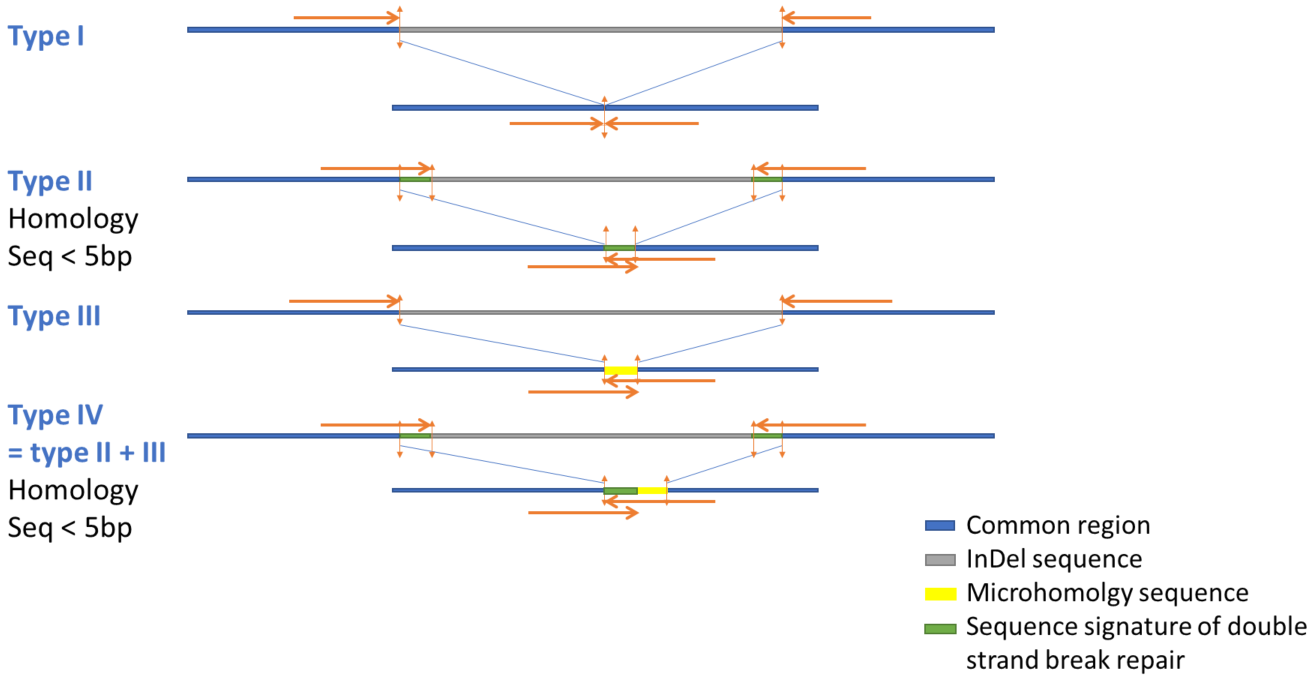

*Figure S3: Schematic representation of four different breakpoint types identified by PINDEL at InDel breakpoints according to the presence of micro-homology sequence or not in place of the deleted sequence.*

*Horizontal arrows represented the forward and reverse BP probes designed at breakpoint site.*

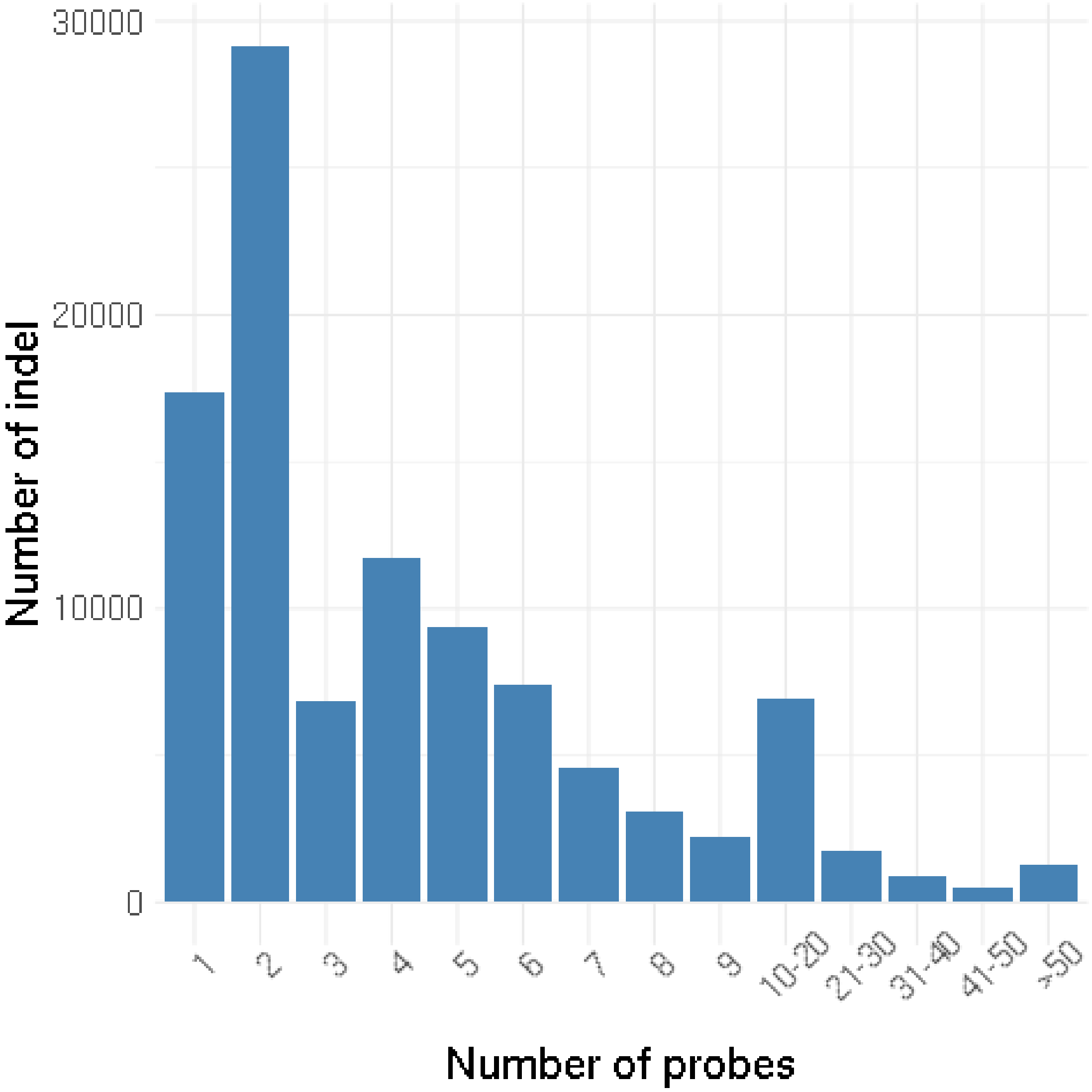

*Figure S4: Distribution of probe number per InDel for 105,927 InDels genotyped with the array.*

*InDels with at least 50 probes were represented in the same category.*

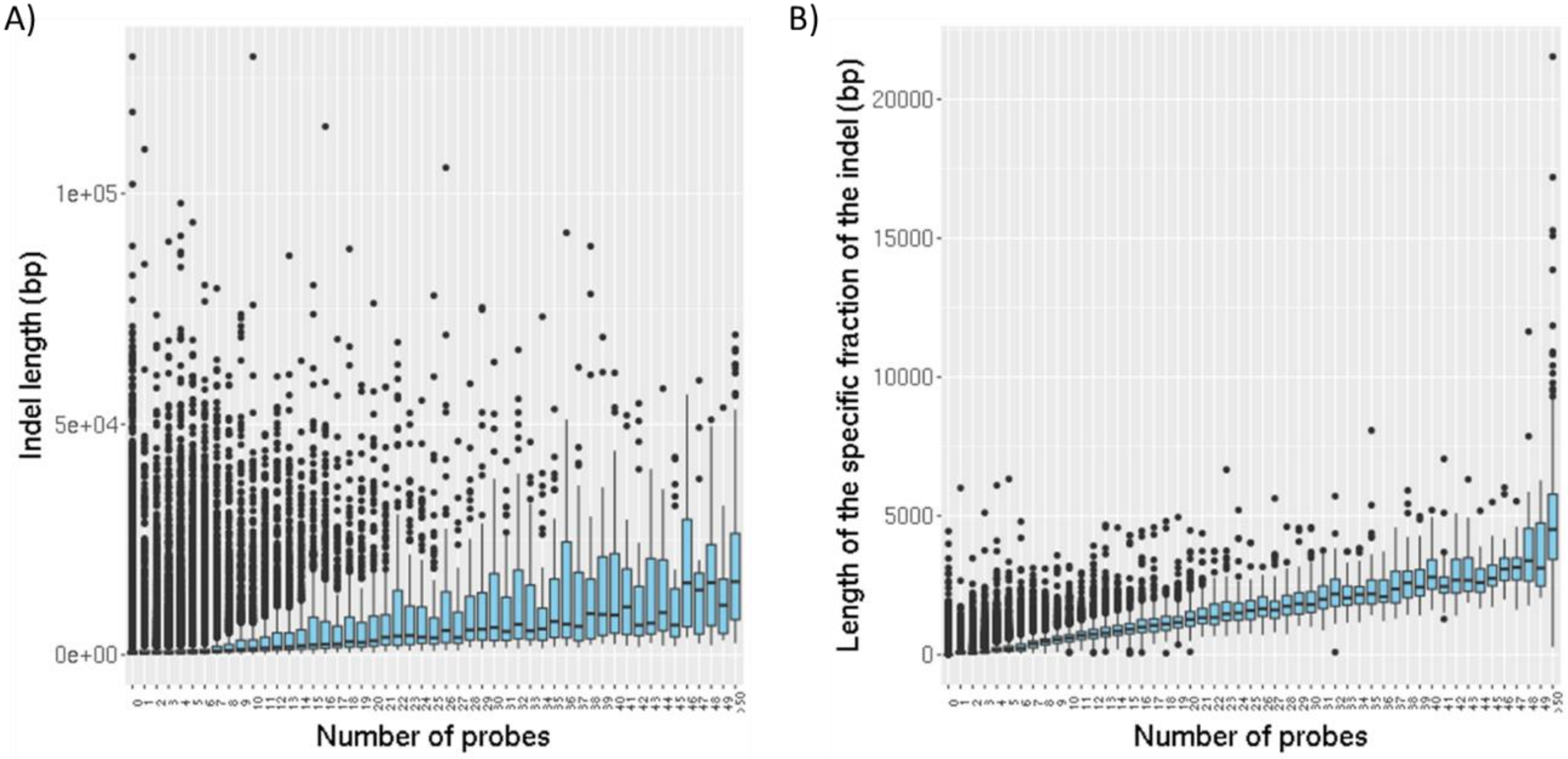

*Figure S5: Relationship between probes number genotyping the InDel and A) the InDel length B) cumulated length of specific sequence (PARs) within InDel.*

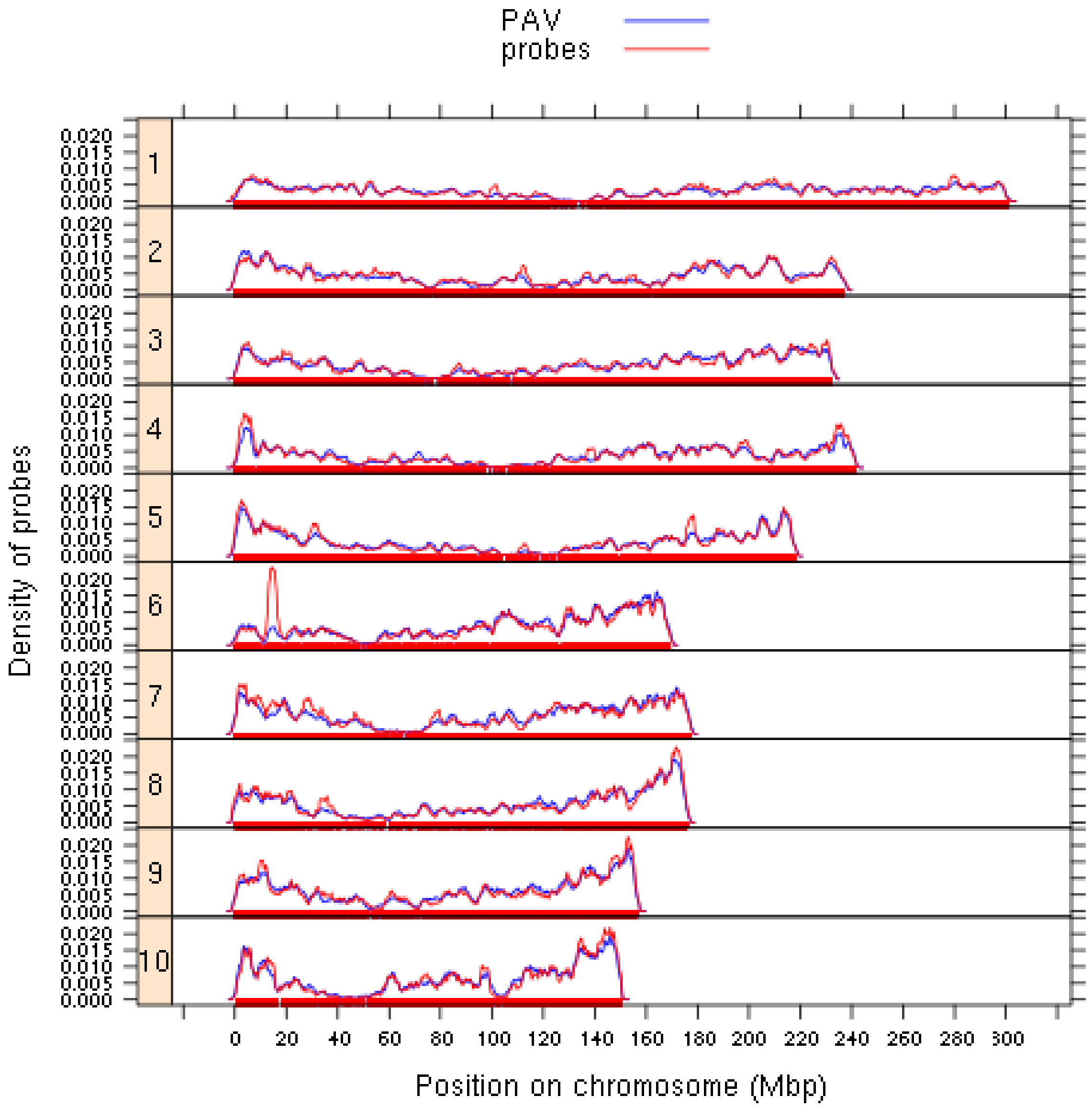

*Figure S6: Variation of probes and InDels density across the 10 maize chromosomes.*

*Red and blue lines represented the density of 237,257 probes and 43,117 InDels anchored in the maize genome, respectively.*

A)

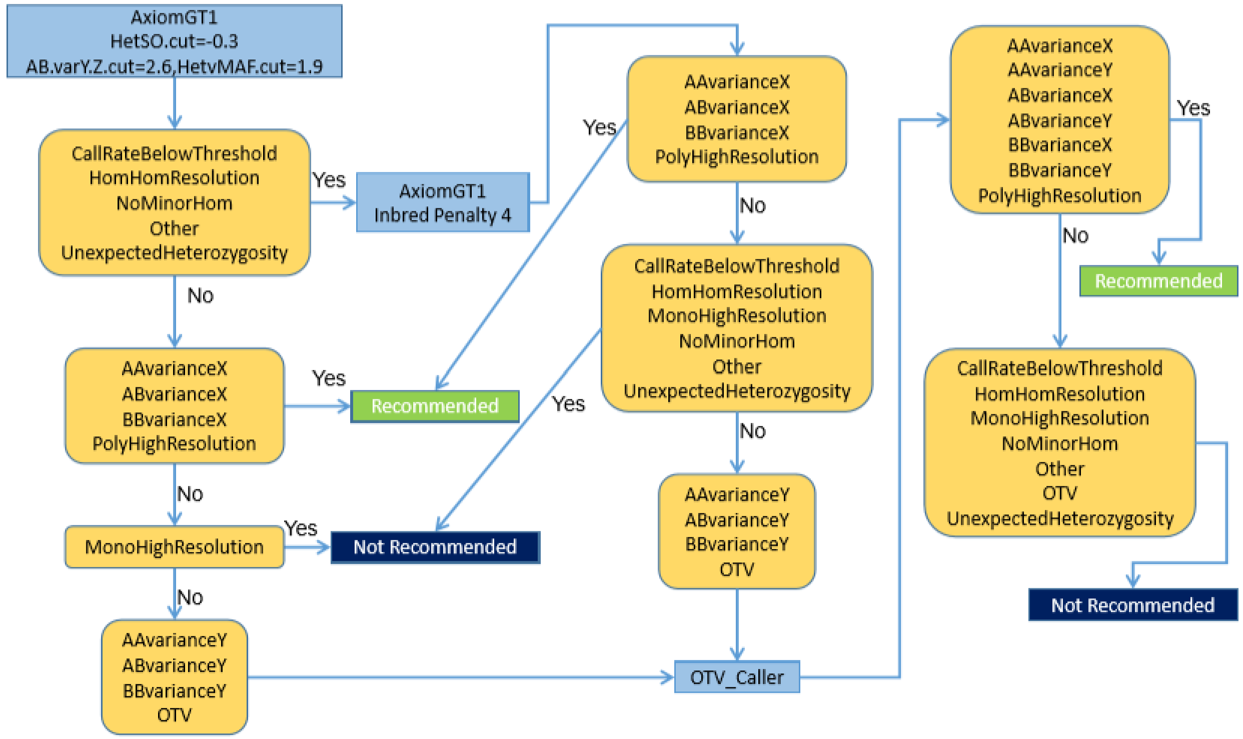
AxiomGT1
HetSO.cut=-0.3
AB.varY.Z.cut=2.6,HetvMAF.cut=1.9
CallRateBelowThreshold
HomHomResolution
NoMinorHom
Other
UnexpectedHeterozygosity
Yes
AxiomGT1
Inbred Penalty 4
No
AAvarianceX
ABvarianceX
BBvarianceX
PolyHighResolution
Yes
Recommended
No
MonoHighResolution
Yes
Not Recommended
No
AAvarianceY
ABvarianceY
BBvarianceY
OTV
Yes
AAvarianceX
ABvarianceX
BBvarianceX
PolyHighResolution
No
CallRateBelowThreshold
HomHomResolution
MonoHighResolution
NoMinorHom
Other
UnexpectedHeterozygosity
Yes
No
AAvarianceY
ABvarianceY
BBvarianceY
OTV
OTV_Caller
AAvarianceX
AAvarianceY
ABvarianceX
ABvarianceY
BBvarianceX
BBvarianceY
PolyHighResolution
Yes
Recommended
No
CallRateBelowThreshold
HomHomResolution
MonoHighResolution
NoMinorHom
Other
OTV
UnexpectedHeterozygosity
Not Recommended

B)

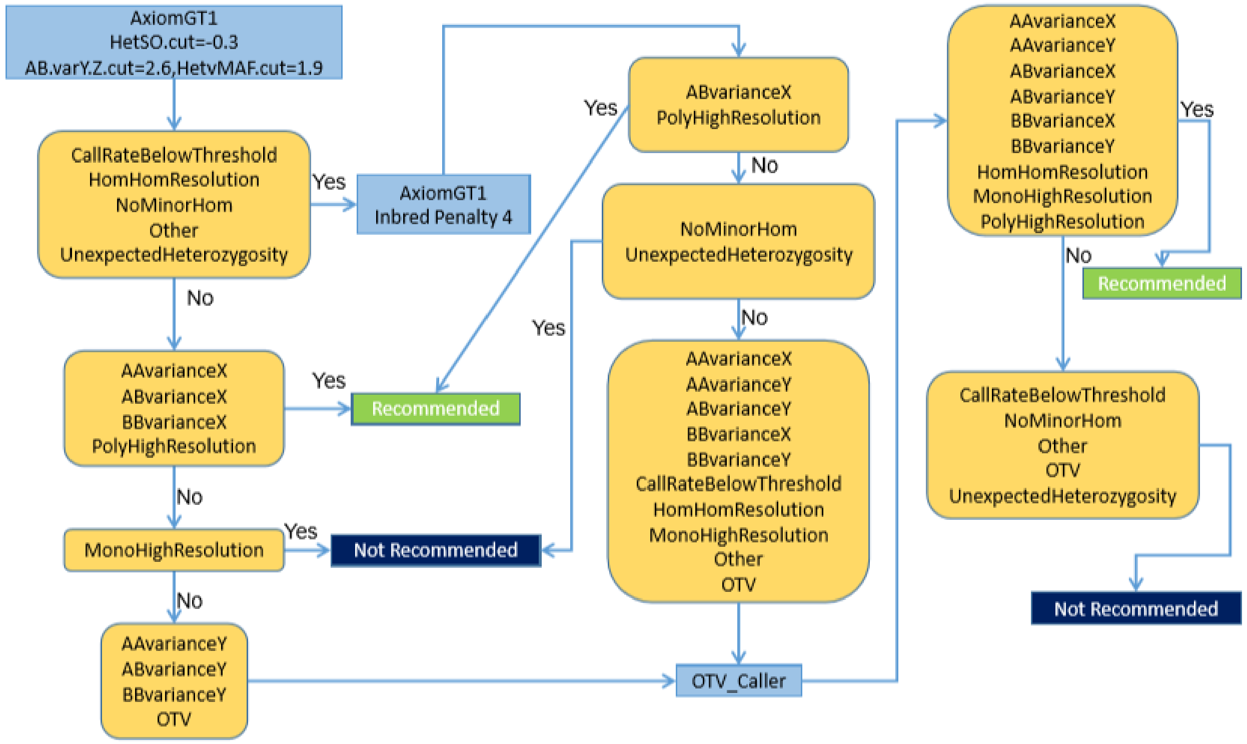
AxiomGT1
HetSO.cut=-0.3
AB.varY.Z.cut=2.6,HetvMAF.cut=1.9
CallRateBelowThreshold
HomHomResolution
NoMinorHom
Other
UnexpectedHeterozygosity
Yes
AxiomGT1
Inbred Penalty 4
No
AAvarianceX
ABvarianceX
BBvarianceX
PolyHighResolution
Yes
Recommended
No
MonoHighResolution
Yes
Not Recommended
No
AAvarianceY
ABvarianceY
BBvarianceY
OTV
Yes
ABvarianceX
PolyHighResolution
No
NoMinorHom
UnexpectedHeterozygosity
Yes
No
AAvarianceX
AAvarianceY
ABvarianceY
BBvarianceX
BBvarianceY
CallRateBelowThreshold
HomHomResolution
MonoHighResolution
Other
OTV
OTV_Caller
AAvarianceX
AAvarianceY
ABvarianceX
ABvarianceY
BBvarianceX
BBvarianceY
HomHomResolution
MonoHighResolution
PolyHighResolution
Yes
No
Recommended
CallRateBelowThreshold
NoMinorHom
Other
OTV
UnexpectedHeterozygosity
Not Recommended

C)

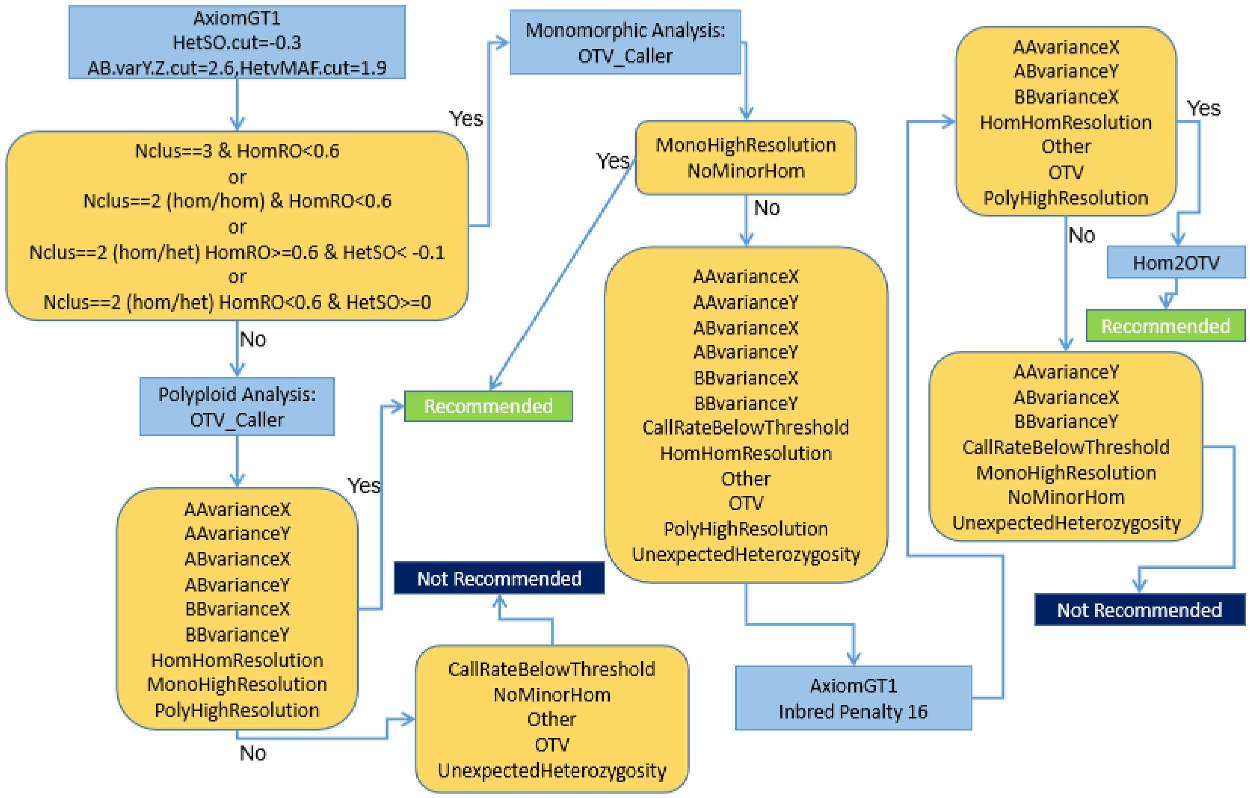

*Figure S7: Three dedicated Affymetrix pipelines used for calling InDel polymorphisms from the fluorescent intensity variation of BP probes (A), OTV probes (B) and MONO probes (C).*

*Each probe was classified into different categories according to the number of clusters, the call rate, and quality metrics of the clustering based on the position, variance, and separation of different clusters. In order to retrieve the best clustering for each probe, successive steps of clustering using different clustering algorithms (light blue square, Axiom GT1, OTV caller, Hom2OTV) or/and with different parameters. According to their classification at each step (yellow square) and threshold used for quality metrics, probes could be classified as recommended (green square), not recommended (blue square), or to be submitted to another step. According to the probe type (BP, OTV), algorithms and parameters of these different steps of clustering varied. For instance, BP and OTV did not differ for the first step of clustering but differed for the following step, since more categories were called with "OTV caller" for OTV probes. Besides this small difference, BP and OTV probes were called in the same way (A, B). A new algorithm (Hom2OTV) was specifically developed for calling InDel genotypes from MONO probes (C) since we expected only 2 clusters (absence / presence) that varied exclusively for fluorescent intensity (Size/Y-axis) rather than for fluorescent intensity ratio (Contrast/X-axis) between two labelled nucleotides. At the end, all probes were classified into 14 categories, which translated to either recommended or not recommended. HetSo: The Heterozygous Cluster Strength Offset  measured the difference in the signal intensities of the genotype clusters as the heterozygous cluster should have higher signal intensity on average compared to the homozygous clusters.  HomSo: The Homozygote Ratio Offset" displayed the distance of the homozygous clusters to Contrast (X-axis) equals 0, to detect potentially misplaced clusters. Nclus: The number of cluster*

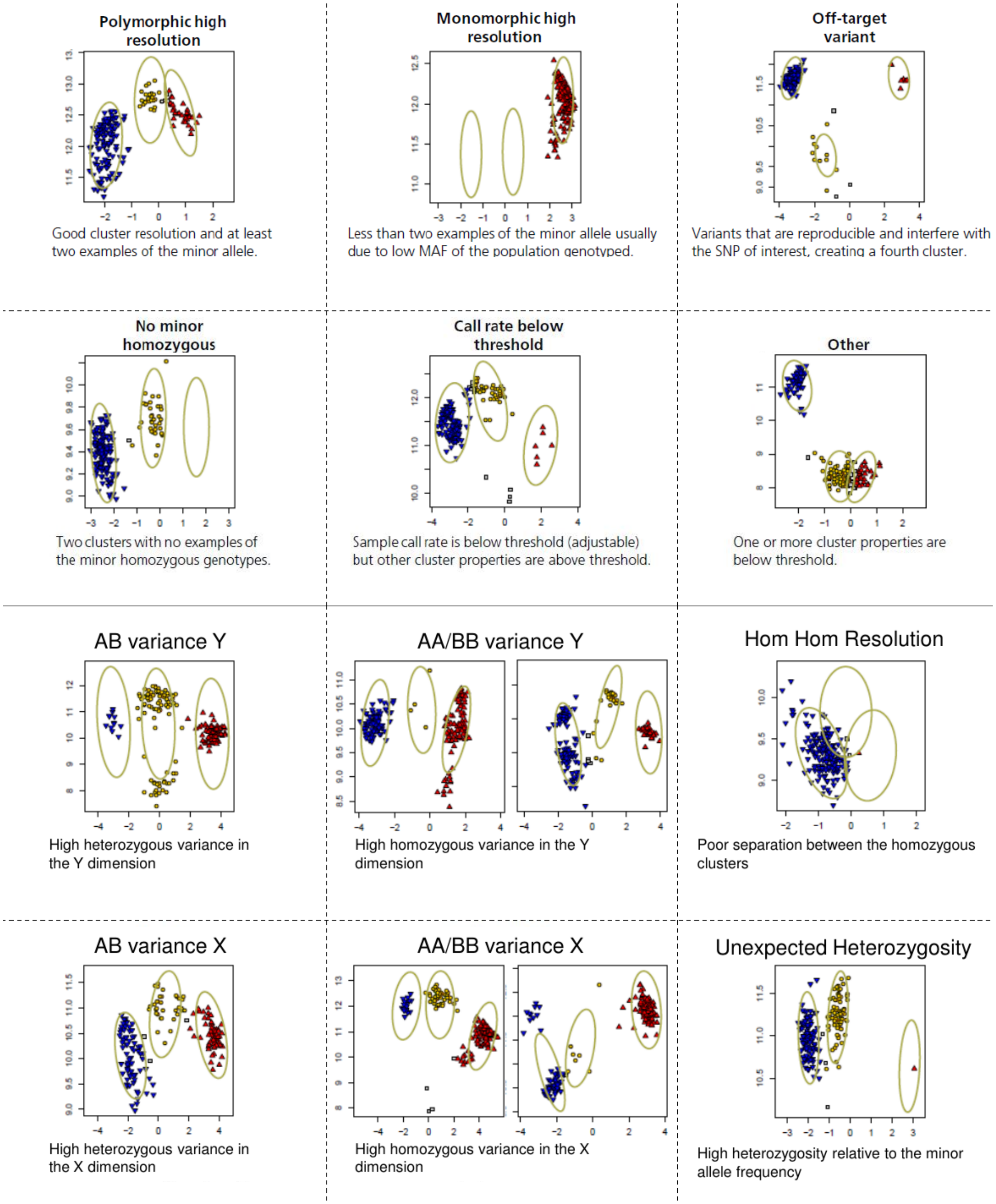

*Figure S8: Example of clustering based on probes fluorescence (intensity in y-axis and contrast in x-axis), for 14 different classifications of probes assigned by the Affymetrix® algorithm.*

*Classifications are based on cluster number, separation of cluster, variance of fluorescent intensity, contrast within clusters, and call rate. For each classification, particular characteristics is described under the figure.*

A)

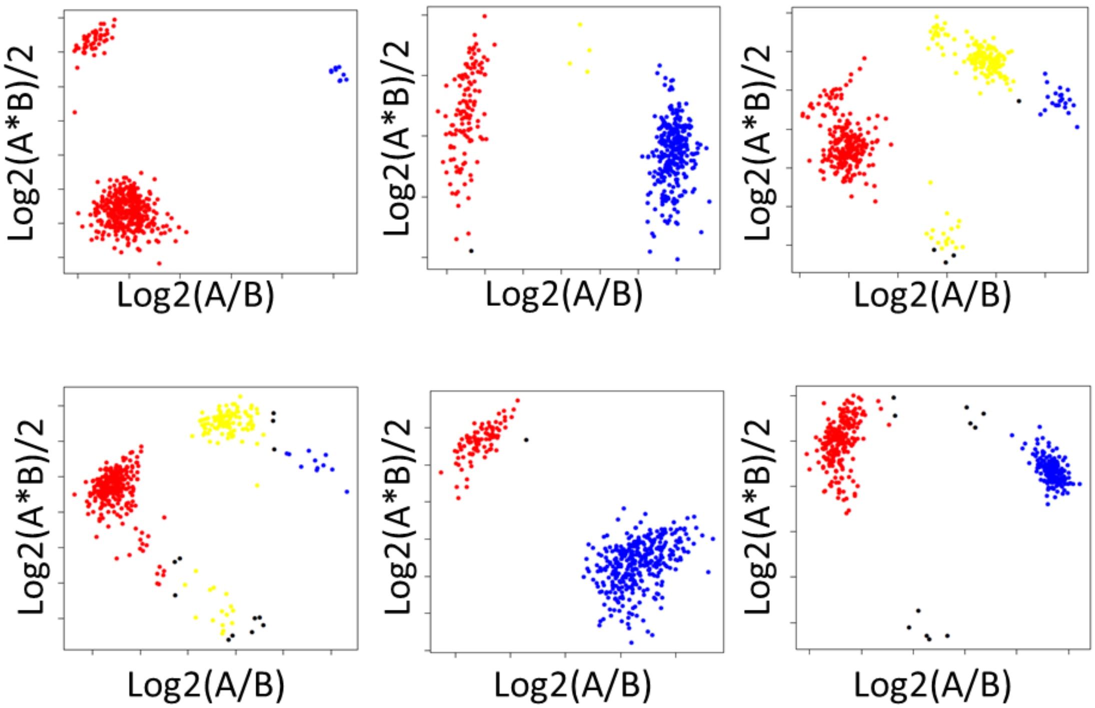

B)

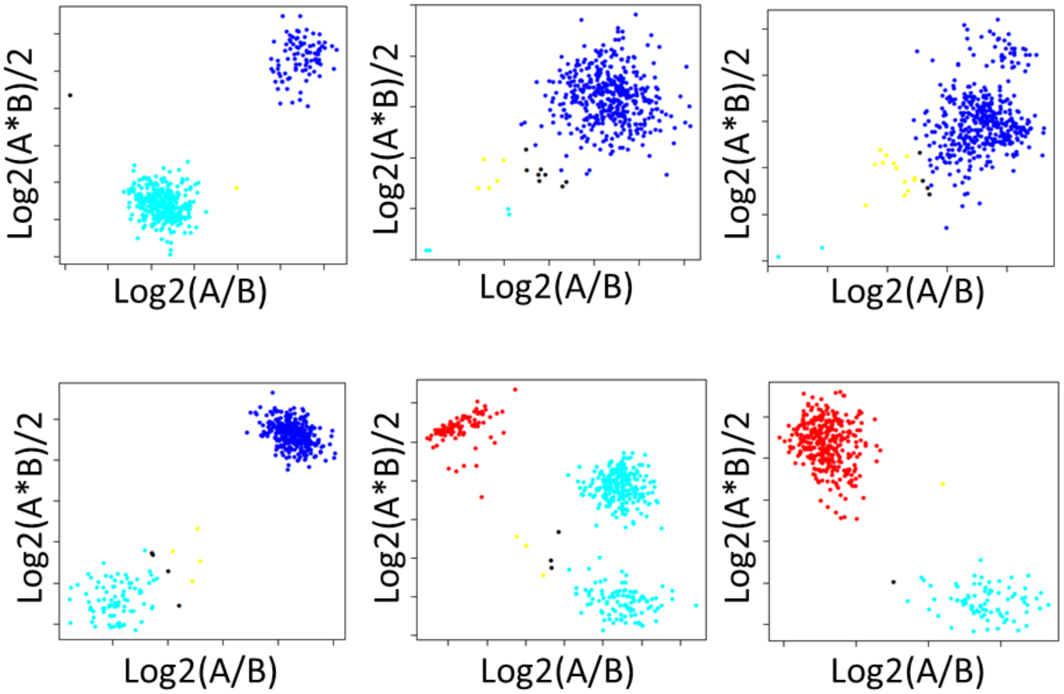

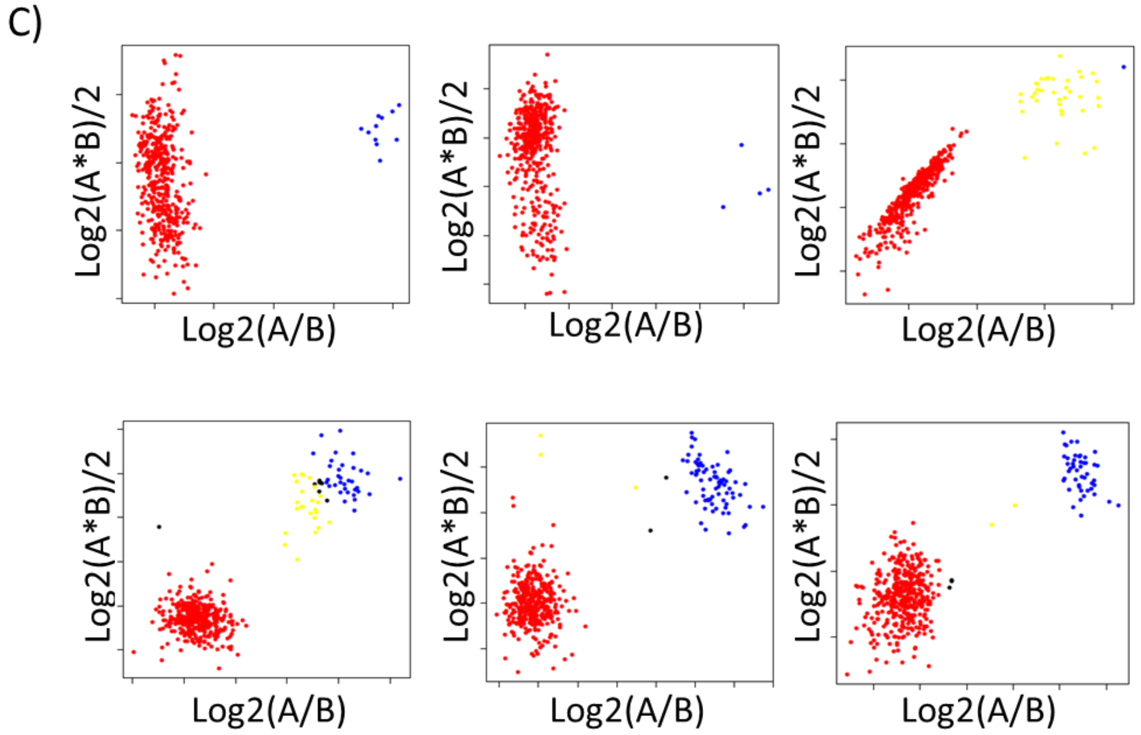

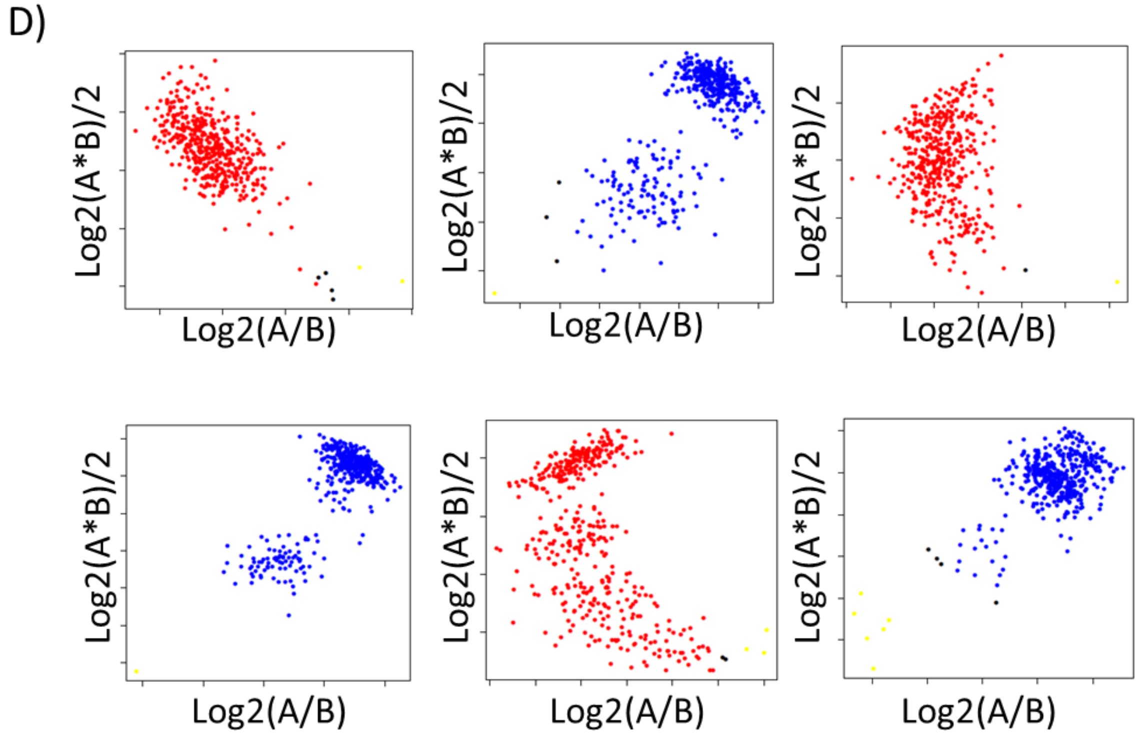

*Figure S9: Example of clustering for 6 randomly probes in different classifications.*

*A) 17,562 OTVs classified as SNP, B) 68,562 MONOs probes with an unexpected heterozygous cluster, C) 1,981 MONOs classified as SNP, and D) 9,525 MONO probes with an unexpected heterozygous cluster but without cluster for absence of the sequence. Blue, red and yellow dots indicated that the sequence were present and were homozygous for allele A (AA) allele B (BB) or heterozygous (AB). Cyan dots indicated that the probes did not hybridize and that the sequence of probes was therefore absent.*

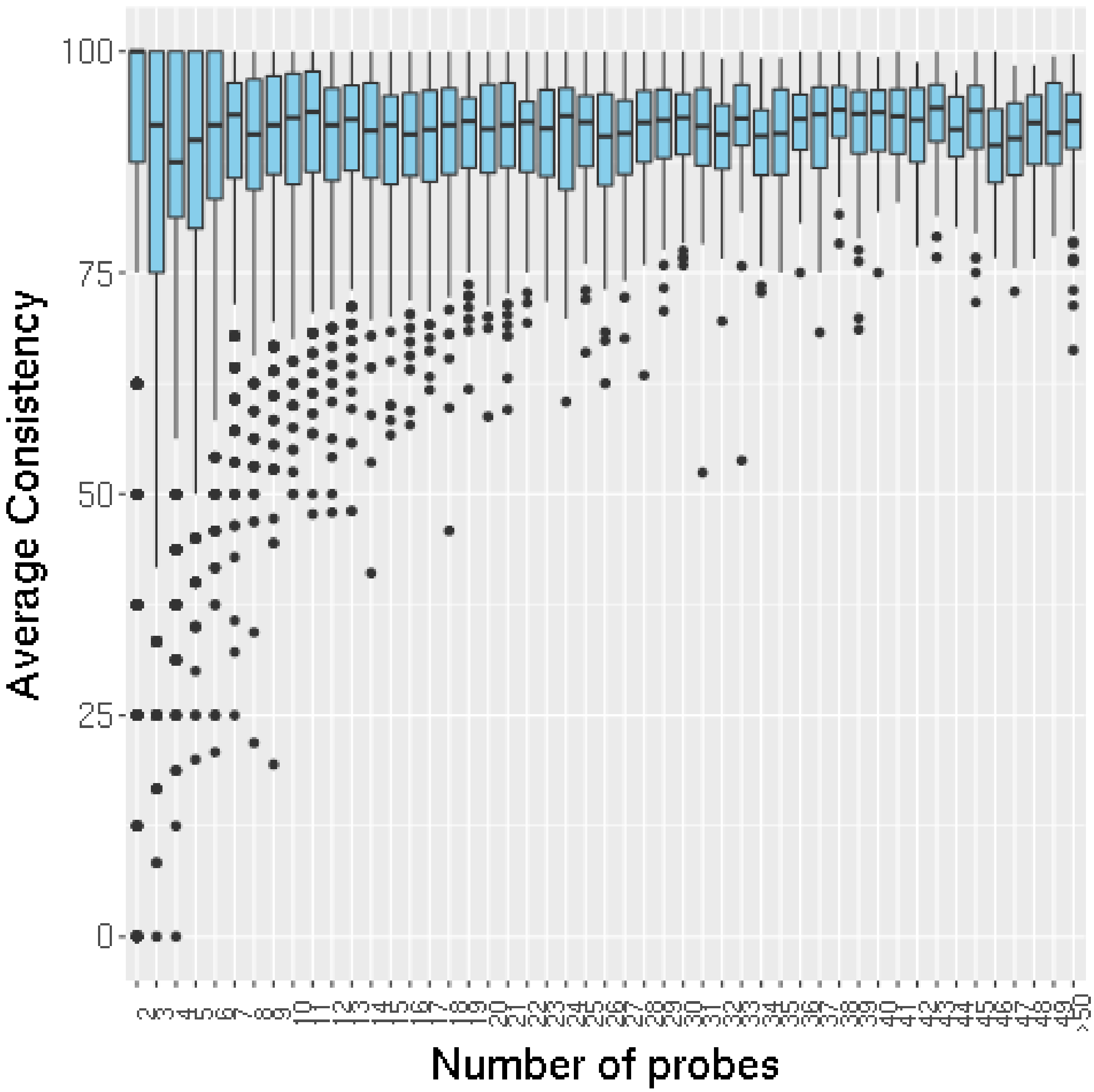

*Figure S10: Variation of the distribution of the average consistency rate (%) of InDels between expected and observed genotyping of probes according to number of probes within the InDel.*

*Only InDels genotyped with at least two probes were considered, and InDels with more than 50 probes were classified in one category (>50).*

53

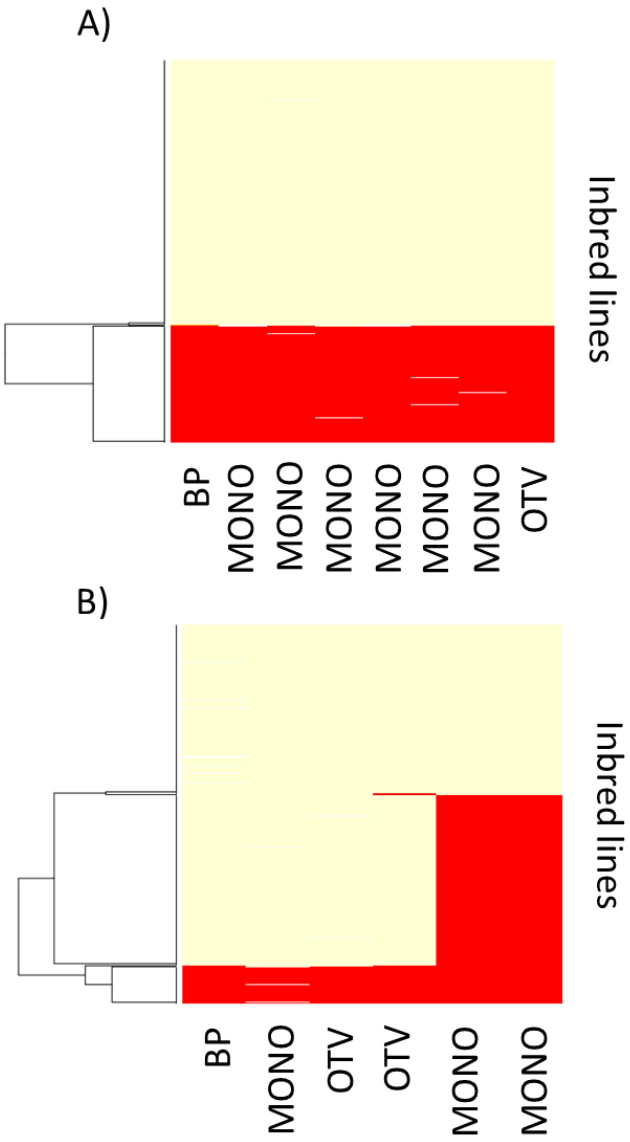

*Figure S11: Haplotype of two InDels genotyped with multiple probes (in column) for 362 individuals (in rows).*

*A) InDel CNVMAIZE_DEL_12046 genotyped with 8 probes displayed two haplotypes indicating either that sequence was totally present (yellow) or absent (red) in 362 individuals. B) InDel CNVMAIZE_DEL_16880 genotyped with 6 probes displayed 3 different haplotypes according to probes genotyping indicating that sequence was only partially absent in some individuals. Inbred lines were ordered by hierarchical clustering according to their similarity based on the genotyping of individual probes. Yellow and red indicated that the sequence of the probes was either present or absent, respectively*

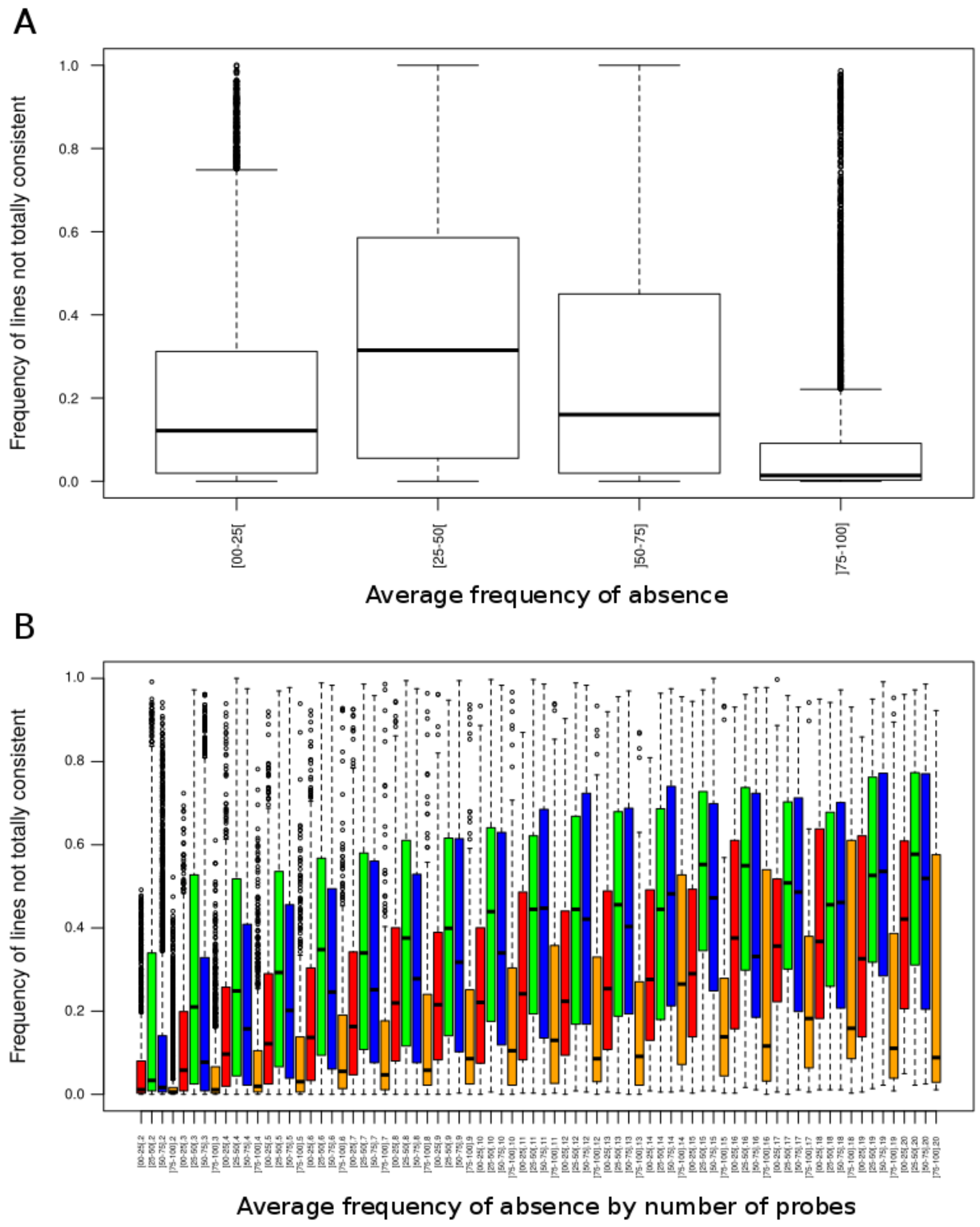

*Figure S12: Effect of average frequency of absence across 362 lines on consistencies between probes genotyping within InDels.*

*A) Boxplot of frequency of lines not fully consistent according to average frequency of absence across 362 lines B) Boxplot of frequency of lines not fully consistent according to number of probes and average frequency of absence across 362 lines. Each InDel are categorized in four classes according to their average allelic frequency of absence: 0-0.25 (red in pane B), 0.25-0.50 (green in pane B),0.50-0.75 (blue in pane B), 0.75-1 (orange in pane B).*

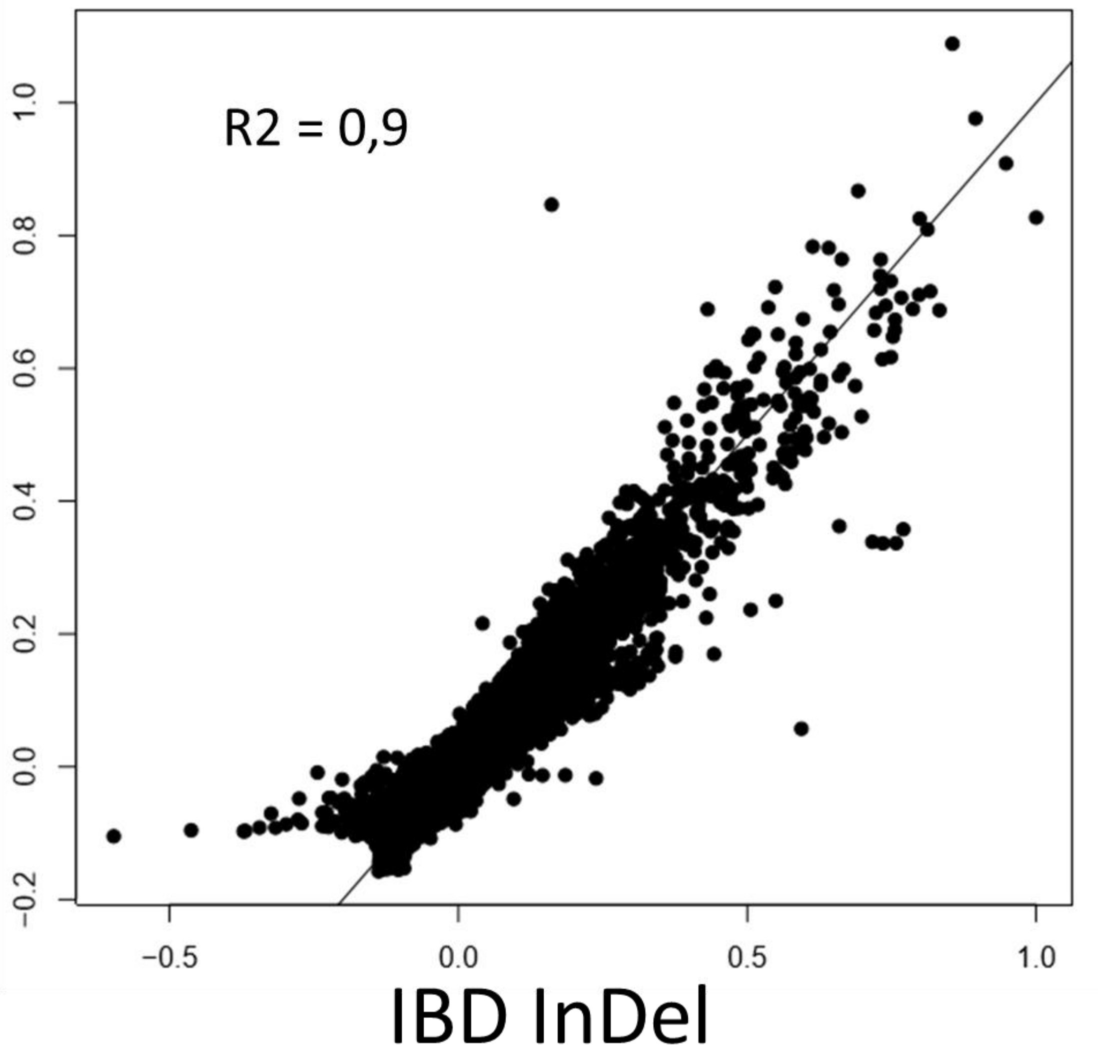

*Figure S13: Comparison of kinship between 362 inbred lines estimated with 57,824 InDels and with 28,143 SNPs from the 50K Illumina genotyping array.*

*Solid lines represent bisectrix.*

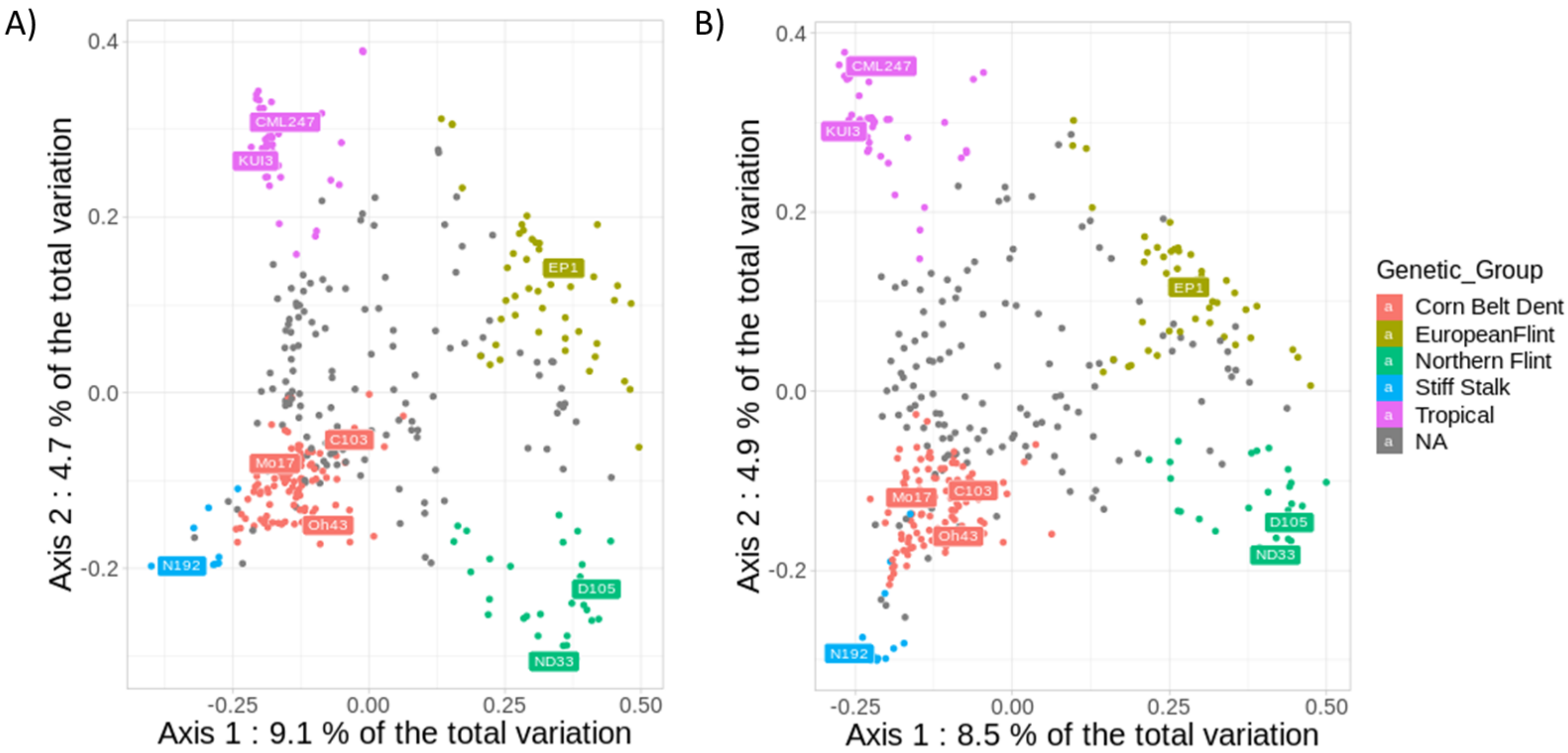

*Figure S14: Principal coordinate analysis on the genetic distance between 360 inbred lines from an association panel (B73 and F2 were excluded) estimated by A) 57,824 InDels and B) 28,143 SNPs.*

*Colors represented the assignation of the inbred lines to the 5 genetic groups defined by admixture using Panzea SNPs from the 50K Illumina array, when the probability of assignment to a group (membership) was greater than 60%. Inbred lines that are not assigned to a group (membership<60%) were considered admixed and colored in gray. Common name of maize accessions typical of each genetic group were indicated.*
